# Toward A Whole-Brain Interface Enabled by High-Resolution Surface Electrophysiology

**DOI:** 10.64898/2026.09.24.754261

**Authors:** Elton Ho, Katrina Barth, Jacqueline Dister, Aneil Srivastava, Adam J. Poole, Federico Claudi, Molly V. Lucas, Joshua S. Miller, Mark Hettick, Yoon Woo Byun, Ramin Anushiravani, Mark Murphy, Peter Konrad, Sanjay Bhatia, Geoffrey Adams, Steven Carte, Dominique Ottonello, Joshua Bederson, Ignacio Saez, Christopher Kellner, Gelana Tostaeva, Brian H. Kopell, Daniel D. Cummins, Sedat Ulkatan, Saadi Ghatan, Iahn Cajigas, Bijan Pesaran, Pedro G. Borges, Qasim Qureshi, Jarl Haggerty, Zhengjia Wang, Nathan Crone, Chetan Bettegowda, Kurt Lehner, Steve Luo, Miguel Angrick, Rebecca Greene, Youssef Comair, Joshua P. Aronson, Michael Lawton, Prashanth Prakash, Tianhao Lei, Joon Hei Lee, Marc W. Slutzky, Matthew Tate, Angelique Paulk, Gavin Dunn, Benjamin Grannan, Mijail D. Serruya, Kevin Hines, Chengyuan Wu, Robert Rosenwasser, Assaf Berger, Elad I. Levy, Adnan H. Siddiqui, Jonathan Riley, Jason M. Davies, Jayme Strauss, Michael J. Mager, Craig H. Mermel, Benjamin I. Rapoport

## Abstract

The goal of building a whole-brain interface—capable of generating a real-time digital representation of the state of the entire central nervous system—has long been framed as a hardware challenge of recording simultaneously from ever-larger numbers of microelectrodes. Advances in high-resolution neural interfaces now enable electrophysiologic interaction with the human nervous system at spatial and temporal resolutions previously unattainable without disrupting tissue architecture or physiology. These capabilities create new opportunities for mechanistic neuroscience, neuroprosthetics, and precision-guided neurosurgery. With the advent of modern microelectrode array technology, however, the whole-brain interface has also become a data challenge. Rather than pursuing large-scale human brain recording purely through a single device placed in a single patient, we demonstrate here that systematic multi-subject aggregation, across brain regions and behavioral paradigms, in a common anatomical framework, can achieve something functionally similar to a whole-brain interface.

Here we present an approach to a functional atlas of the human central nervous system derived from high-resolution surface electrophysiology recordings obtained in the first 70 patients implanted with an anatomically conformal, thin-film neural interface positioned in more than 100 locations. The atlas comprises >100,000 electrode deployments and catalogs placement across a broad range of neuroanatomic targets spanning the motor, sensory, language, auditory, and associative domains, as well as brainstem and spinal cord structures that have historically been difficult to access at high electrophysiologic resolution. Across participants, aggregated recordings reveal somatotopic organization, including digit representations that follow a lateral-to-medial ordering consistent with classical homuncular maps, with transitions between adjacent representations occurring over spatial scales of approximately 0.8–2.0 mm; canonical frequency-band signatures, such as movement-related beta-band suppression and focal high-gamma (70–200 Hz) increases during motor and speech behaviors; and task-modulated population dynamics in both evoked and spontaneous activity across cortical, brainstem, and spinal cord regions. These findings yield insight into patterns of mesoscale electrophysiological dynamics conserved across the entire human central nervous system, suggesting that fine-scale functional organization is preserved across individuals when aligned to common anatomical landmarks. We further demonstrate several applications of these high-resolution recordings, including real-time neural decoding from upper extremity motor cortex to enable real-time neuroprosthetic control, and from ventrolateral cortex to decode cortical correlates of spoken language, facial movements, and complex facial expressions of emotion, supporting initial feasibility of combined verbal and nonverbal communication interfaces.

Together, these results establish a paradigm for approaching large-scale interfaces with the human brain, as well as a starting point for a scalable, extensible atlas of human electrophysiologic function designed to accelerate discovery across basic neuroscience, translational research, and clinical neurotechnology.

## Introduction

A central goal in applied neuroscience is to link human neural activity to sensation, movement, language, and cognition with sufficient resolution to reveal mechanisms while remaining compatible with routine use in human patients. Noninvasive approaches such as functional magnetic resonance imaging (fMRI) ^1^, magnetoencephalography (MEG) ^2^, and electroencephalography (EEG) ^3^ provide whole-brain coverage but are limited by spatiotemporal resolution and indirectness of measurement, whereas intracranial electrophysiology provides direct access to neural population dynamics with millisecond precision ^4^ but has historically been constrained by sparse spatial sampling and heterogeneous, case-specific coverage. As a result, widely accessible human brain atlases are principally anatomical ^5^ or imaging-based ^6^; even with the availability of multi-center databases of intracranial electrophysiologic recordings ^7–9^, and after the advent of multi-center functional-imaging-based parcellation ^10^ ^11^, there is still no widely used, multi-subject, high-resolution electrophysiologic reference atlas spanning multiple functional domains and extending beyond the cerebral cortex ^12^.

Recent advances in microfabrication, materials science, and surgical technique have begun to shift this landscape by enabling high-density surface electrophysiology with substantially finer spatial sampling ^13^ than standard clinical electrocorticography ^14^. Conformable thin-film arrays ^13^ can be placed transiently on exposed neural surfaces ^15^, enabling recordings at submillimeter pitch without penetrating tissue and, in some implementations, allowing reversible placement in intraoperative or short-term extraoperative settings. These developments create an opportunity to treat human electrophysiology not only as a per-patient mapping tool but also as a set of scalable measurement techniques capable of supporting systematic aggregation across subjects, tasks, and anatomical targets.

High-resolution surface recordings occupy a regime that is complementary to existing modalities. Penetrating microelectrodes can capture unit activity at cellular scale but disrupt tissue ^16^ and, while enabling high-resolution recordings in selected settings ^16^ ^17^ , are challenging to deploy broadly across clinical contexts and distributed anatomical targets. Standard clinical electrocorticography (ECoG) grids, in contrast, are safe and widely used but sample coarsely ^18^ ^19^, blurring steep functional gradients and limiting the ability to resolve mesoscale organization and population dynamics that may be essential for decoding and mechanistic inference ^20^ ^21^; this limited resolution applies as well to clinical stereo-electroencephalography (sEEG). High-density surface electrophysiology offers the possibility of bridging this gap ^13^ ^15^ ^21^ ^22^ by combining millisecond temporal fidelity with spatial sampling dense enough to reveal mesoscale structures, while remaining compatible with the temporary exposures that occur routinely in neurosurgical care as well as with emerging long-term implants.

Most intracranial electrophysiology datasets also remain anatomically narrow. The overwhelming majority of high-quality human recordings come from cortical surfaces or depth electrodes placed for epilepsy evaluation ^12^, leaving brainstem and spinal cord physiology comparatively underrepresented in modern, high-resolution datasets ^23^ ^24^ ^25^ ^26^ despite their central role in sensorimotor integration, cranial nerve function, and autonomic control. Yet, in select surgical contexts, these regions become safely accessible for brief intervals. A surface-based recording approach that is anatomically conformal and clinically practical makes it feasible to capture electrophysiologic signatures from these “frontier” structures and integrate them into a unified coordinate framework.

Here we lay the groundwork for a functional atlas of the human central nervous system derived from high-resolution surface electrophysiology in the first 70 patients implanted with an anatomically conformal, thin-film neural interface ^15^. Across these cases, the atlas aggregates data from 87,342 electrodes spanning (1) primary sensorimotor cortex, (2) ventrolateral frontal and superior temporal regions supporting speech production and auditory or receptive-language processing, (3) lateral prefrontal cortex engaged during decision-making paradigms, and (4) surgically accessible brainstem and spinal cord surfaces, including dorsal column and trigeminal complex regions and thoracic spinal cord sites. Recordings were obtained across intraoperative and extraoperative settings, with standardized task sets and stimulus-locked paradigms when feasible, and with consistent preprocessing and quality metrics to support cross-subject comparison.

We designed the atlas to serve two goals: first, as a scientific report of high-resolution human electrophysiology across multiple functional systems, and second, as a methods and data resource that can be extended as enrollment grows. To support mechanistic inference and translation, we focus on reproducible, interpretable signatures, including somatotopic gradients and phase reversals around the central sulcus, canonical spectral modulations during movement and speech, stimulus-locked evoked responses in cortex and in ascending sensory pathways, and task-modulated population dynamics ^27^ ^28^ ^29^ ^18^ ^30^. We further include demonstrations of decoding to illustrate how dense surface recordings can support applications that require high spatial-frequency features often attenuated in lower-resolution recordings.

By combining dense surface electrophysiology, multi-subject registration, and standardized metadata and code, this work establishes a scalable framework for building an evolving electrophysiologic atlas of the human central nervous system. We anticipate that the atlas will enable cross-subject synthesis of fine-scale functional organization, benchmarking of neural decoding models, and integration with structural and functional imaging, accelerating discovery across basic neuroscience, translational neurotechnology, and precision-guided neurosurgery.

Elements of the work presented here, including the system configuration ^15^, methods of human intraoperative use ^31^ (including burr-hole ^32^, intra-sulcal ^33^, and cisternal ^34^ electrode array deployment; as well as concurrent use of multiple arrays ^35^), central sulcus mapping ^33^, and motor and speech decoding ^32^ ^36^ ^37^, are described in depth parallel manuscripts. In this manuscript, we first present a multi-subject overview of electrode placement and data composition across the cohort of patients studied, establishing the spatial extent, resolution, and uniformity of recordings (Fig. 1). We then assess reproducibility, anatomical alignment, and cross-subject variability across domains, establishing a framework for aggregation and reuse of high-resolution human electrophysiologic data (Fig. 2). We then characterize high-resolution functional organization along the Rolandic cortex, including phase reversals and fine-scale somatotopy revealed by peripheral stimulation and movement paradigms (Fig. 3), followed by population-level motor representations during upper-extremity tasks (Fig. 4). We next describe language-, speech-, (Fig. 5) and facial-expression–related dynamics (Fig. 6) in ventrolateral frontal and superior temporal regions, highlighting distinct yet overlapping micro-scale patterns associated with symbolic and emotive behaviors, and auditory responses and tonotopic organization within auditory-associated temporal cortex (Fig. 7). Extending beyond the cerebral cortex, we demonstrate surface electrophysiology from surgically accessible brainstem and spinal cord sites, resolving stimulus-locked activity and spatial organization within ascending sensory pathways and cranial nerve territories (Fig. 8). Together, these results define an initial functional atlas of the human central nervous system and provide a foundation for its continued expansion.

**Figure 1:**
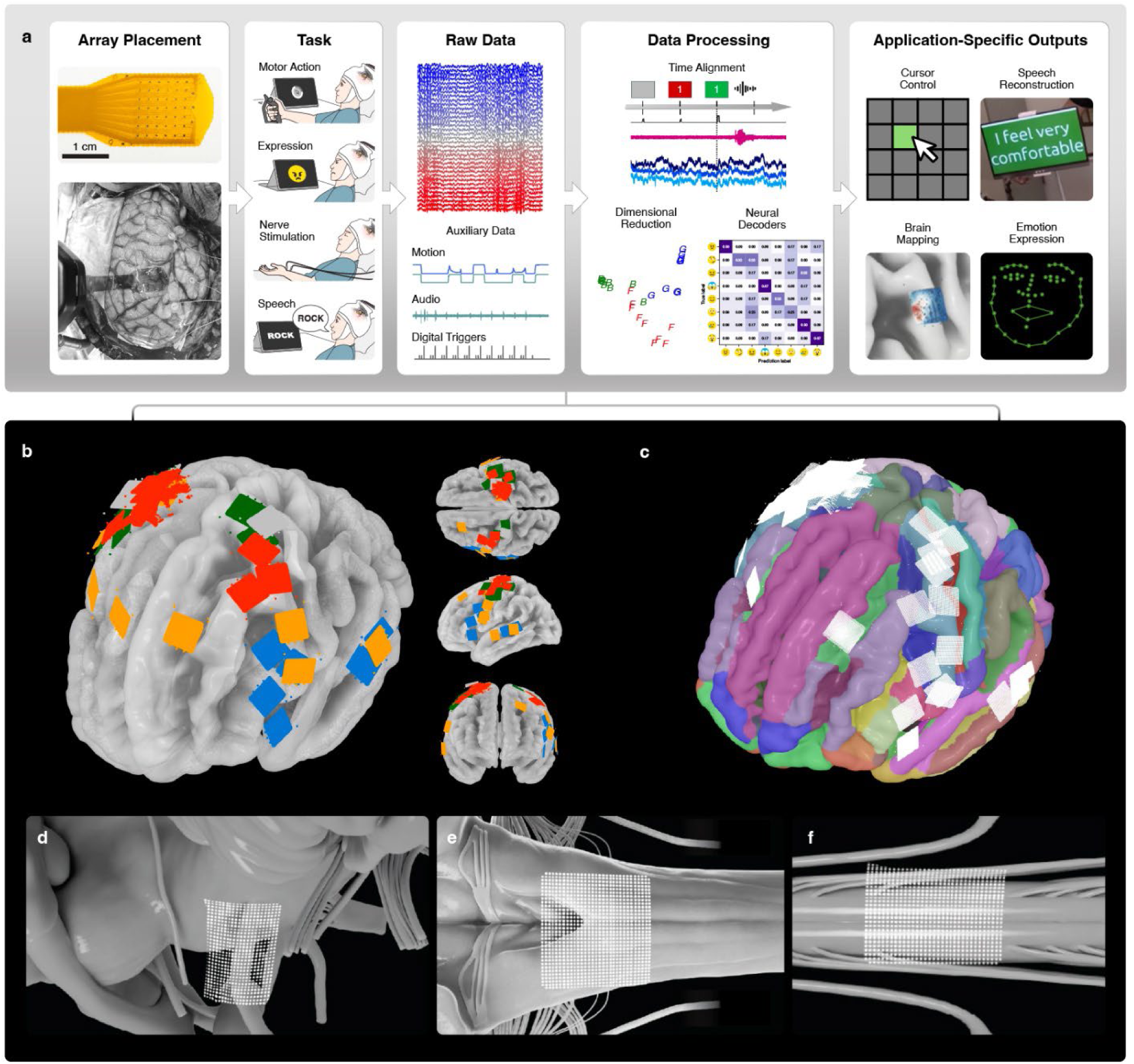
Atlas overview and multi-subject electrode coverage. (a) Workflow of generation of high-resolution neural data atlas and associated annotations to support task-specific use cases. Neural data and annotations (including motion, audio, and event triggers, as appropriate) were collected while patients were executing tasks with arrays placed over accessible cortical areas. The collected data were preprocessed to extract desirable neural features (high-gamma amplitude, for example), and then further processed to extract information and outputs useful for brain-computer interface function or other electrophysiologic tasks. (b) Composite 3D rendering of a template brain (fsaverage) overlaid with electrode placements from all patients. Placements are color-coded by patient task: hand and finger movement (motor and somatosensory) targeting primary sensorimotor cortex (red); facial and speech expression targeting face motor and ventrolateral precentral and inferior frontal opercular areas (blue); central sulcus localization targeting the Rolandic region (green); and other applications (orange), including, but limited to, risk-reward assessment, auditory response, cortico-cortical responses, and device characterization. (c) Visualization of the same arrays with the brain template color-coded according to the Destrieux atlas. (d-f) 3D rendering of arrays placed over the trigeminal nerve, brainstem, and thoracic spinal cord, respectively, with left being rostral.

**Figure 2.**
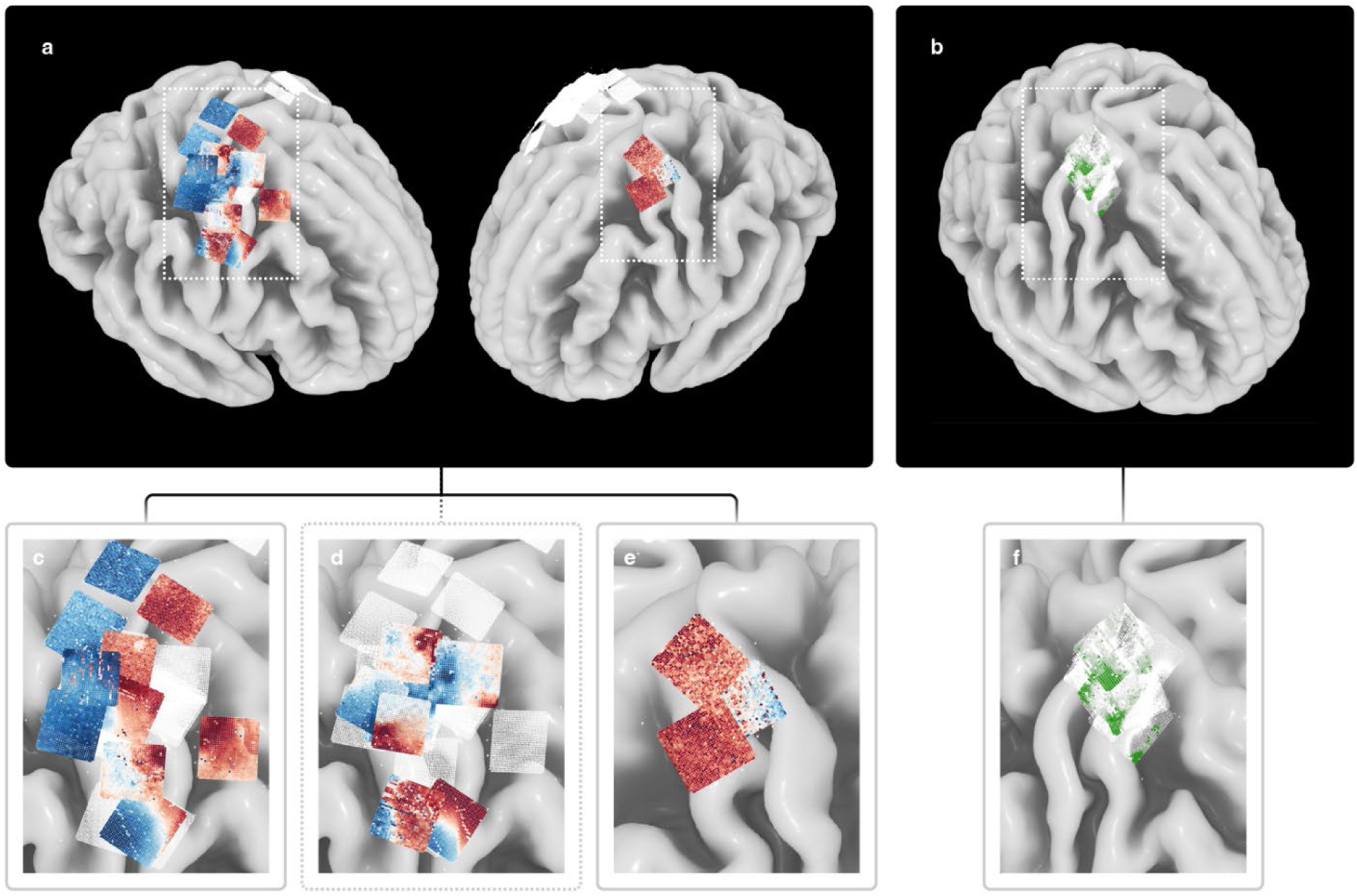
Consistency and variability across subjects in the high-resolution neural atlas. (a) Spatial voltage maps of median nerve somatosensory evoked potentials (SSEPs) from a subset of 12 subjects, representing 21 deployment locations, co-registered to a common anatomical reference frame, demonstrating a reproducible N20/P20 polarity reversal aligned to the central sulcus. Red indicates positive voltage, blue indicates negative. (b) Group-level maps of high-gamma (70–200 Hz) activity during hand gesturing, showing convergence of task-related activity onto the precentral hand knob region across subjects after anatomical alignment. (c) Ten arrays (color-graded arrays), on the right hemisphere, showing alignment between the functional boundary and the central sulcus. Whited-out arrays correspond to arrays presented in (d). (d) Eight additional arrays, presented separately for visual clarity. Some arrays represent functional boundaries less well-aligned to the central sulcus, potentially due to physiological distortions from underlying pathology (including mass lesions). Whited-out arrays correspond to arrays presented in (c). (e) Three arrays on the left hemisphere. (f) Zoomed view of high-gamma maps from (b).

**Figure 3:**
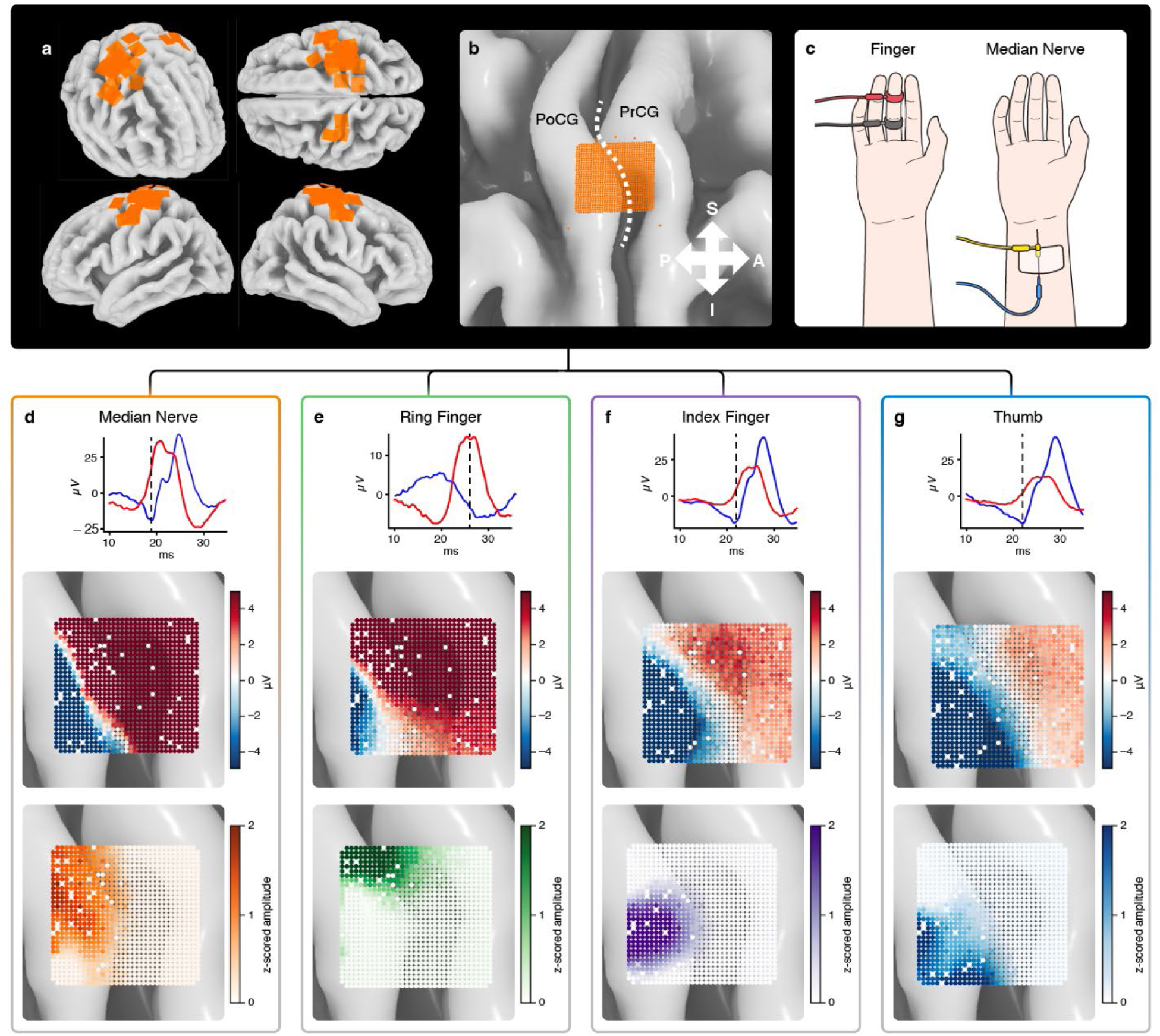
Sensorimotor mapping across the Rolandic cortex. (a) A 3D rendering of 21,462 electrodes deployed through 21 arrays in a subset of 13 patients. Arrays typically straddled the central sulcus when one array was deployed, while for cases in which multiple arrays were deployed, electrode arrays may be completely anterior or posterior to the central sulcus. (b) Zoomed-in view of an example array placed over the central sulcus (dashed line), for which we collected data from upper extremity stimulation. (c) Diagram of the median nerve and finger-level stimulation via circular ring electrode stimulation. (d) Trial-averaged SSEP responses from stimulation of the median nerve. (e-g) SSEPs for three individual fingers. Each top panel shows the trial-averaged voltage traces (filtered from 30–1000 Hz) from a negative channel (blue) and a positive channel (red). The dashed line indicates the time point post-stimulus at which a negative peak (N20) was identified over the somatosensory cortex. Middle panels show the voltage at each electrode over the array at the indicated time point. The lower panels show the z-score normalized peak-to-peak amplitude of the SSEP responses at each electrode over the array, highlighting the area with greatest response amplitude for each stimulus location.

**Figure 4:**
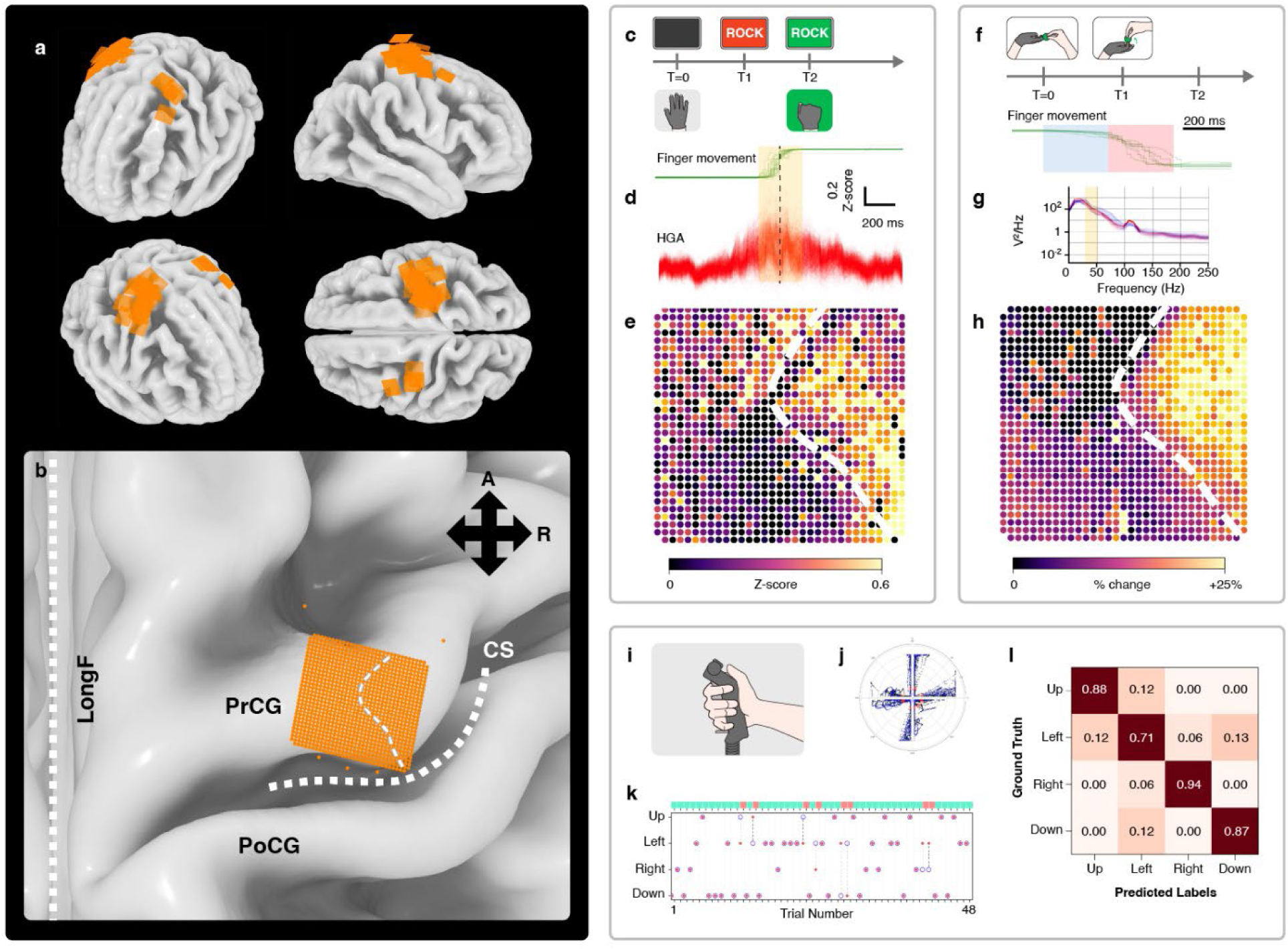
Upper extremity motor representations resolved with high-density cortical arrays. (a) Composite 3D rendering showing electrode placements targeting primary motor cortex across all subjects included in this analysis. (b) Zoomed view of an example array positioned over the precentral hand knob region. To guide the eye, the dashed line over the precentral gyrus indicates the empirically identified boundary between regions with and without strong event-related high gamma activity, reflected in other subpanels. (c) Experimental paradigm for active hand gesturing. The green lines each represent a trial of finger flexion. (d) Trigger-averaged high gamma amplitude. Each red line represents one channel, averaged over 60 trials and time-aligned to movement onset, as indicated by the dashed line. (e) High-gamma spatial map of a temporal snapshot during peak finger velocity. (f) Experimental paradigm for passive finger movement. The green lines each represent the finger position of a trial of finger extension. (g) Power spectral density of neural activity in the time windows as indicated by the blue (early motion) and red (peak motion) time windows in (f). (h) Spatial map of increase in low gamma band, as a ratio between peak versus early motion, highlighted as the yellow band in (g). (i-j) Center-out joystick movement, with example trajectories in polar representation. (k) Trial-by-trial classifier output (red dots) compared to ground truth (blue circles). The cyan and red trials indicate correct and wrong predictions, respectively. (l) Confusion matrix for classifying four movement directions for a single subject.

**Figure 5:**
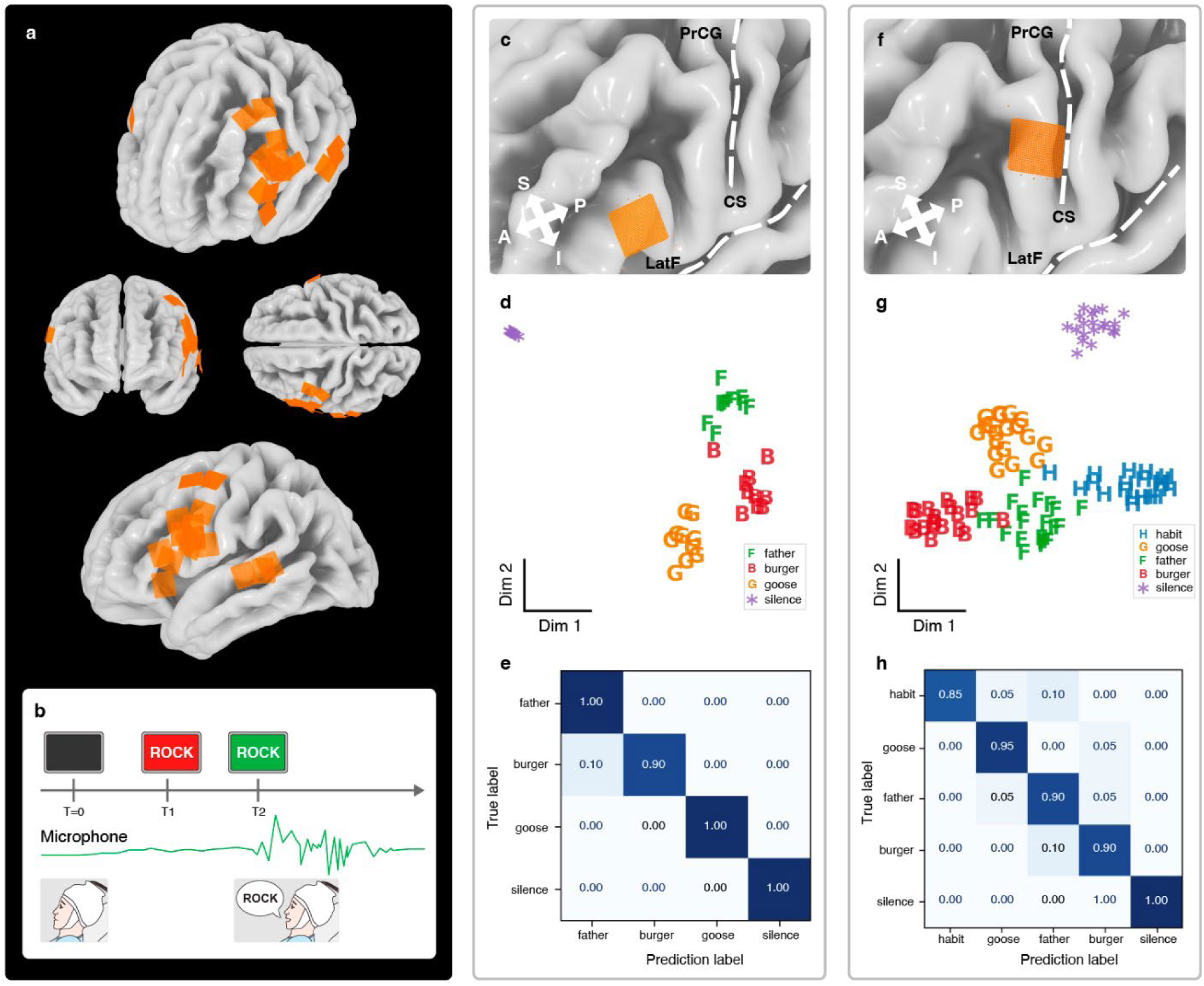
Speech neural activity in ventrolateral and motor cortex. (a) Composite three-dimensional rendering showing electrode placements over ventrolateral frontal and temporal cortex across all subjects included in this analysis. (b) Experimental paradigm for word utterance tasks. (c) Zoomed view of an example array positioned over inferior frontal gyrus. (d) Dimensional reduction (t-SNE) of neural data corresponding to each speech trial. Uttered words: *F* = “Father”, *B* = “Burger”, *G* = “Goose”, and * = Silence. (e) Confusion matrix for 4-class decoding. (f) Zoomed view of an example array positioned over precentral gyrus. (g, h) Dimensional reduction (t-SNE) and confusion matrix with an additional uttered word *H* = “Habit”.

**Figure 6:**
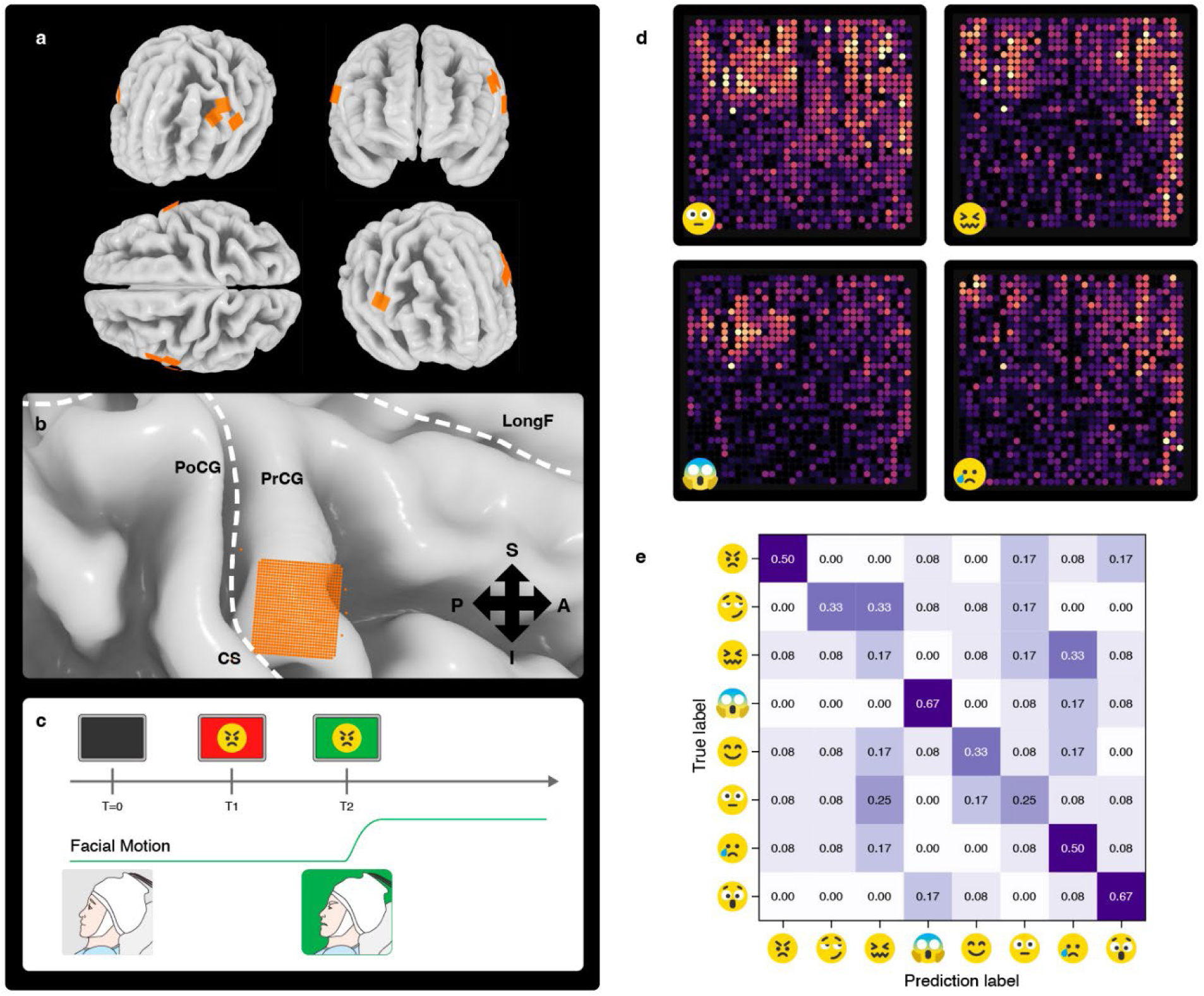
Facial expression neural activity in ventrolateral and motor cortex. (a) Composite three-dimensional rendering showing electrode placements over ventrolateral prefrontal and motor cortex across 4 of the 8 subjects in whom neural correlates of facial expression were studied. (b) Zoomed view of an example array positioned over precentral gyrus. (c) Experimental paradigm for facial expression tasks. (d) Spatial map of high-gamma activity corresponding to four distinct facial expressions (“Neutral,” “Frustrated,” “Terrified,” “Sad”). Note the spatially distinct cluster of active electrodes for each emotion. (e) Confusion matrix for an 8-class facial expression decoder. The diagonal represents the classification accuracy for each expression, showing separability between distinct emotive states, including “Angry,” “Smirking,” “Frustrated,” “Terrified,” “Happy,” “Sad,” “Shocked,” and “Neutral.”

**Figure 7:**
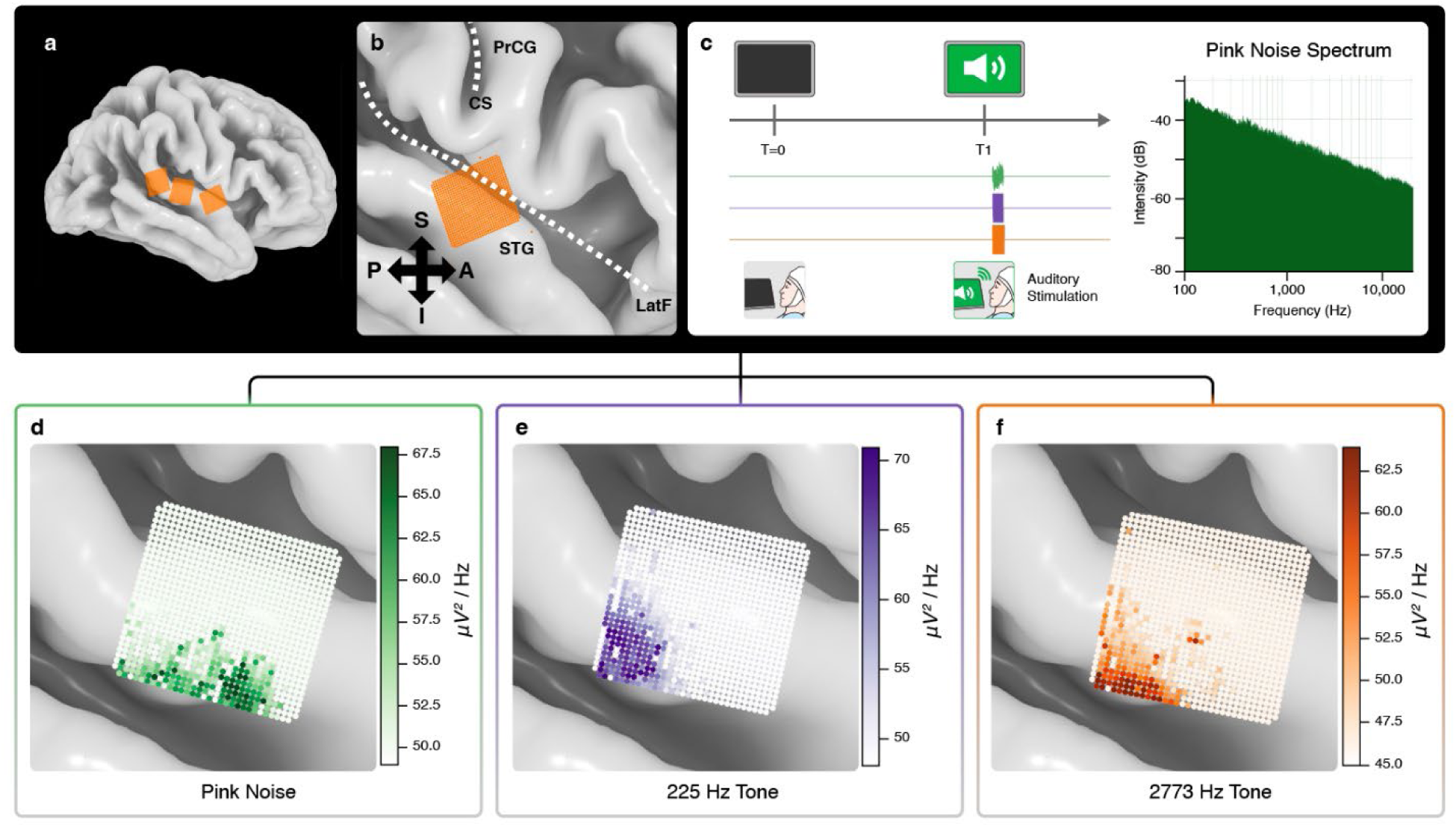
High-resolution mapping of auditory stimuli. (a) Composite three-dimensional rendering showing electrode placements over the superior temporal gyrus and auditory-associated cortex across three subjects. (b) Zoomed view of an example array positioned over the superior temporal gyrus (STG). (c) Experimental paradigm for passive listening tasks, depicting the timeline of audio presentation. (d-f) Spatial maps of trial-averaged (N=50) neural signal power in the 9.8–34.2 Hz range during the 1 second following presentation of pink noise, 225 Hz pure tones, and 2773 Hz pure tones, respectively. Note the distinct, spatially separated distributions of activity for low-versus high-frequency stimuli, consistent with tonotopic organization.

**Figure 8:**
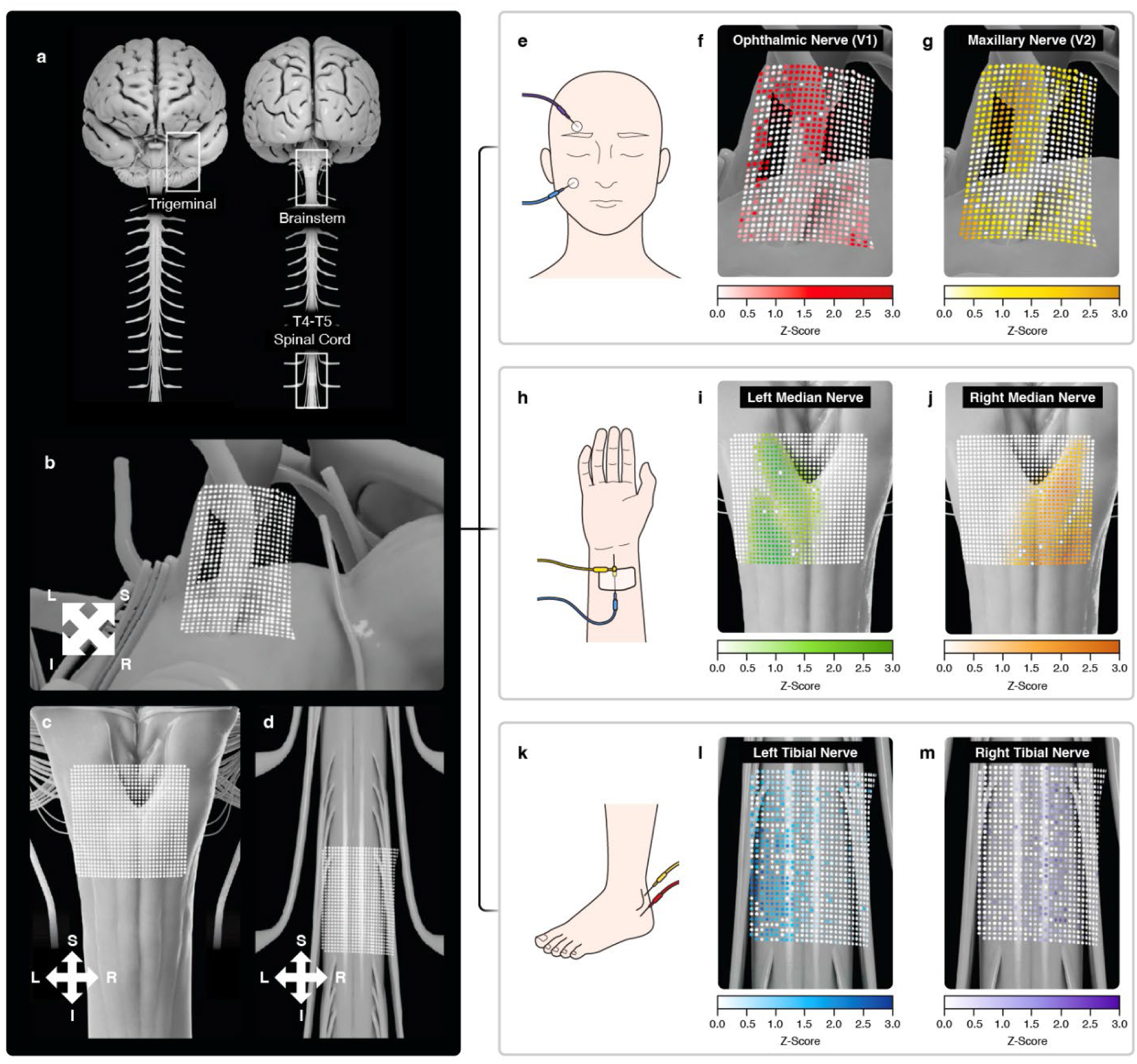
High-resolution surface electrophysiology of the brainstem and spinal cord. (a) Composite three-dimensional renderings showing electrode placements over the trigeminal root entry zone, brainstem, and spinal cord surfaces. (b-d) Zoomed views of example arrays positioned over the trigeminal root entry zone, brainstem near the cervicomedullary junction and ventral pons, and thoracic spinal cord, respectively. (e-g) Spatial maps of trial-averaged, normalized evoked signal amplitude during stimulation of the ophthalmic (V1) and maxillary (V2) branches of the trigeminal nerve. (h–j) Spatial maps of trial-averaged normalized evoked signal amplitudes recorded over the brainstem surface in the region of the dorsal columns during left and right median nerve stimulation. (k-m) Normalized evoked signal amplitudes recorded over the T4–T5 dorsal spinal cord during left and right tibial nerve stimulation. All maps depict task-aligned evoked responses projected onto array geometry.

## Results

### 1. Descriptive Statistics of the High-Resolution Neural Data

The aggregated neural data described here comprises high-resolution electrophysiological recordings from 70 patients already undergoing neurosurgical procedures. Data were collected across 86 arrays deployed on the cortex, brainstem, spinal cord, and cranial nerves, totaling 87,342 micro-electrode channels recording from the central nervous system, at 103,598 unique microanatomic locations (Fig. 1; Table 1). Arrays were implanted temporarily, and recordings were performed in intraoperative settings either awake or under general anesthesia, as well as in extraoperative contexts during the postoperative hospital stay, depending on clinical indication and experimental paradigm.

**Table 1:** Summary of electrode array deployments and recorded data across anatomical targets in the neural data atlas. For each anatomic location, the table reports the number of patients and deployed arrays, patient state during recording (degree of anesthesia), primary experimental paradigm, total number of electrodes in contact with neural tissue, and total volume of raw neural data collected (GB). The number of deployed arrays includes instances in which the same physical array was repositioned (redeployed) to enable recording in a different anatomic location. Combined totals across all deployments are shown in the final rows. Across the cohort, some electrode arrays were repositioned once or twice (N=15) to sample from multiple anatomic regions during a single operative session (resulting in a total of 102 locations mapped), while some cases had multiple arrays deployed (N=14). Consequently, when categorized by target location, the aggregate totals for patients (N=83) and arrays (N=102) exceed the actual number of unique patients (N=70) and physical arrays (N=86) utilized in the study.

| Anatomic Location | Number of Patients | Arrays Deployed | Awake/Asleep | Paradigm | Number of Electrodes | Data (GB) |
| --- | --- | --- | --- | --- | --- | --- |
| Superior temporal gyrus | 7 | 9 | Both | Auditory stimuli or speech decoding | 8,962 | 156 |
| Inferior frontal gyrus | 4 | 4 | Awake | Speech decoding | 4,054 | 203 |
| Middle frontal gyrus | 1 | 1 | Awake | Hand motor decoding | 960 | 143 |
| Superior frontal gyrus | 4 | 5 | Awake | Speech decoding and cognitive responses | 5,120 | 158 |
| Central sulcus | 15 | 22 | Both | Mapping somatosensory responses or motor decoding | 22,422 | 1,151 |
| Precentral gyrus (upper extremity motor) | 25 | 29 | Awake | Motor decoding or mapping somatosensory responses | 29,440 | 7,506 |
| Precentral gyrus (face motor) | 10 | 12 | Both | Speech or facial expression decoding or face stimuli | 12,160 | 1,327 |
| Postcentral gyrus | 8 | 10 | Both | Hand sensorimotor or facial expression decoding | 10,240 | 1,457 |
| Trigeminal nerve | 3 | 3 | Asleep | Mapping cranial nerve responses | 3,072 | 101 |
| Brainstem | 4 | 5 | Asleep | Mapping sensory responses | 5,120 | 344 |
| Spinal cord | 2 | 2 | Asleep | Mapping peripheral nerve responses | 2,048 | 78 |
| Totals (Unique) | 83 (70) | 102 (86) |  |  | 103,598 (87,342) | 12,624 |

Across all deployments, the hardware demonstrated robust recording performance, with an average functional channel yield of 89.1% ± 12.2% and a mean electrode impedance of 1.17 MΩ ± 0.50 MΩ at 1 kHz (N=69,254 included electrodes from N=77 arrays were assessed intraoperatively prior to data acquisition). Using a 400 µm electrode pitch, the system achieved a spatial sampling density of 625 electrodes cm⁻², exceeding that of standard 10 mm pitch clinical electrocorticography arrays by more than 600-fold ^38^.

Electrode coverage was distributed across a broad set of central nervous system targets (Fig. 1a–b), enabling systematic sampling of multiple functional domains. Cortical placements included high-density recordings from primary motor and somatosensory cortex, superior and inferior precentral gyrus, frontal and temporal association cortices, and the internal banks of the central sulcus. In addition to cortical recordings, the atlas includes electrophysiological data from brainstem and spinal cord surfaces, regions that are rarely represented in human electrophysiology datasets due to the geometric and mechanical constraints of legacy electrode technologies, and the challenges of recording from these highly sensitive regions at high resolutions in an atraumatic manner.

In total, the dataset comprises >26.9 hours of intraoperative and >153.9 hours of extraoperative synchronized neural and behavioral recordings, corresponding to 4.7 TB of intraoperative and 7.9 TB of extraoperative raw neural data (Table 1). Recording paradigms included motor execution and imagery, speech perception and production, cranial and peripheral nerve stimulation, and task-free mapping protocols conducted under anesthesia. For each recording block, the dataset includes raw broadband voltage traces, electrode coordinates co-registered to a template space (fsaverage ^39–42^), and time-aligned behavioral and stimulus annotations, including audio, motion tracking, and event triggers where applicable.

Figure 1 provides an overview of the atlas structure and data modalities. Figure 1a illustrates our standardized data collection workflow for generating consistent task-annotated neural data for a multitude of modalities across target locations and patient populations. Generally, this workflow involves array placement, task presentation, patient action or response, recording of raw neural and annotation data, data processing, and downstream hooks to application-specific outputs. With this workflow, composite multi-subject renderings of electrode placements were constructed across cortical, brainstem, and spinal cord surfaces (Fig. 1b-f).

### 2. Reproducibility and Cross-Subject Alignment of High-Resolution Electro-Anatomic Data

To assess reproducibility and inter-subject alignment of neural features captured by the atlas, we examined convergent electrophysiological signatures across subjects during somatosensory and motor activity, after co-registration to a common anatomical reference frame (Fig. 2) ^43^ ^41^ ^44^.

All participants underwent stereotactic-quality brain MRI, CT, or both prior to surgery. In all cases, this stereotactic-quality imaging facilitated the use of clinical-grade image guidance intraoperatively to co-locate the physiologcial data recorded from the array. Electrode registration was estimated using one of two methods: (1) CT-based localization of radiopaque markers, in cases for which intraoperative or postoperative CT was available (N=24); or (2) via intraoperative optical imaging and expert anatomical alignment, in intraoperative cases without cross-sectional imaging (N=46). With each registration verified on the correct target gyrus, we estimated errors to be comparable to the registration accuracy of clinical-standard image guidance, which was calibrated to a maximum of 2 mm.

Across subjects with arrays spanning the central sulcus, median nerve somatosensory evoked potentials (SSEPs) ^27^ exhibited a highly consistent spatial organization. Median nerve stimulation generates a dipole moment within the fold of the central sulcus, creating measurable positive (P20) and negative (N20) peaks over the primary motor and somatosensory cortices, respectively. To quantify array proximity to the central sulcus, the Euclidean distance from each electrode to the nearest precentral-postcentral boundary in MNI152 ^5^ ^45^ ^46^ space was computed using the Destrieux atlas ^47^. The median electrode-to-boundary distance was 6.4 mm (IQR 4.4-9.6 mm; mean 7.4±4.0 mm) with 64% of electrodes falling within 8 mm of the anatomical boundary. Voltage maps computed at the N20/P20 latency demonstrated a reproducible polarity reversal aligned to the sulcal axis, with the transition localized to adjacent electrodes on opposing banks of the central sulcus (Fig. 2a, c-e). Despite variability in array orientation and cortical folding across individuals, this electrophysiological landmark remained spatially stationary relative to cortical anatomy, providing an anchor point for cross-subject alignment of sensorimotor data. Of the 7,674 total electrodes registered over the precentral gyrus, 77.0% exhibited a P20 peak, consistent with the expected electrophysiologic profile of the motor cortex. Correspondingly, 77.7% of electrodes (N=10,494) registered over the postcentral gyrus exhibited the characteristic N20 peak associated with the primary somatosensory cortex. N=13 array deployments were consistent with the expected polarity reversal across the central sulcus for at least 80% of electrodes. N=8 deployments exhibited greater deviations (>20%) of the functional boundary from the anatomical central sulcus (Fig. 2d), likely attributable to arrays positioned predominantly on one side of the sulcus, weaker stimulation coupling, or tumor-related distortion of the underlying cortical geometry.

Similarly, movement-related activity recorded during hand motion tasks showed convergent spatial organization across patients. High-gamma (70–200 Hz) power increases during hand gesturing consistently localized to the precentral hand knob region, with focal clusters of activity occupying homologous anatomical locations across subjects (Fig. 2b, f). The centroid of the 50 most active electrodes was significantly closer to the canonical hand knob than the centroid of remaining active electrodes (mean difference 1.8 mm, exact permutation test, p = 0.031; Wilcoxon signed-rank p = 0.039, N=7). While the precise spatial extent and peak amplitude of responses varied across individuals, the center of mass of task-related activity was directed towards the hand knob area after anatomical alignment, indicating conserved micro-scale hand motor representations within the primary motor cortex.

Together, these analyses demonstrate that high-resolution electrophysiological features recorded with dense surface arrays exhibit reproducible alignment to cortical anatomy across subjects, while preserving biologically meaningful variability at the sub-millimeter scale. The data set thus supports cross-subject comparisons grounded in stable electrophysiological landmarks, while retaining the fine-grained spatial detail necessary to study individual differences in human neural organization.

### 3. Sensorimotor Mapping of Peri-Rolandic Cortex

High-density arrays (N=22) were deployed over the precentral (BA4) and postcentral (BA3,1,2) gyri in 15 subjects, enabling contiguous sampling of the human sensorimotor cortex across the central sulcus (Fig. 3a-b). The 400 µm electrode pitch provided continuous coverage spanning primary motor cortex (M1; Brodmann area 4 ^48^) and primary somatosensory cortex (S1; Brodmann areas 3b, 1, and 2 ^49^), allowing direct measurement of spatial gradients in evoked cortical activity at sub-millimeter resolution.

Functional delineation of the motor-sensory boundary was assessed using somatosensory evoked potentials (SSEPs) elicited by peripheral nerve stimulation. Electrical stimulation of the median nerve (Fig. 3c) produced robust, time-locked cortical responses when arrays traversed the central sulcus (Fig. 3d). Trigger-averaged signals revealed a characteristic N20/P20 polarity reversal ^27^ across adjacent electrodes along the sulcal axis, consistent with a dipolar source configuration spanning anterior and posterior banks of the central sulcus.

Although physiologic phase reversal is typically identified using linear electrode “strips,” and so correspondingly viewed as occurring at a point, the phase reversal boundary is better understood as a line, the resolution of which is dependent on electrode density in an electrode array contoured to the cortical surface spanning the central sulcus. In cases where arrays straddled the sulcus, the polarity transition occurred over distances as small as 0.8 mm (Fig. 3d), corresponding to two electrode spacings, providing a direct physiological estimate of the spatial extent of the motor–sensory transition zone.

In addition to nerve-level SSEPs, fine-scale somatotopic organization ^50^ ^51^ was examined using digit-specific electrical (N=6 patients) (Fig. 3c) or vibrotactile stimulation ^52^ (N=6 patients). Evoked responses were spatially localized and reproducible across trials, with distinct clusters of electrodes exhibiting maximal peak-to-peak amplitudes for individual digits ^53^ (Fig. 3e-g). Similar fine-scale organization was observed for orofacial representations (see Results Section 6).

Together, these recordings provide a high-resolution physiological map of sensorimotor organization across the peri-Rolandic cortex, capturing both macroscopic somatotopy and micro-scale spatial structure within and across functional representations.

### 4. High-Resolution Upper Extremity Somatotopy in Primary Motor Cortex

High-density arrays (N=31, partially overlapping the cohort described in Section 3) were implanted over primary motor cortex (BA4) and, in some cases, adjacent primary somatosensory cortex (BA3, 1, 2) in 27 patients, targeting the precentral hand knob region ^54^ contralateral to the studied upper extremity (Fig. 4a–b). During active, attempted, or imagined finger and hand movements (including motor imagery paradigms ^55^), neural activity exhibited consistent task-locked modulation across subjects ^56^ ^57^, characterized by beta-band (12–30 Hz) power decreases ^28^ and high-gamma (70–200 Hz) power increases ^29^ localized to subsets of electrodes within the array.

These movement-related modulations were spatially focal, typically confined to regions spanning <10 mm^2^, and were reproducible across trials and behavioral conditions when aligned to cortical anatomy and electrode coordinates (Fig. 4c-e). Both active hand gesturing and passive finger movement (Fig. 4f-h) paradigms elicited structured patterns of gamma activity, with partially overlapping but distinguishable spatial distributions across the array.

Group-level topographic maps revealed conserved somatotopic gradients ^58^ ^32^ ^36^ across the precentral and postcentral gyri, with transitions between adjacent finger representations captured at the spatial scale of the electrode pitch. In addition to finger-level representations, distinct activation patterns corresponding to hand, wrist and forearm movements were observed within the same arrays, with Kruskal’s normalized stress ^59^ in the 0.075-0.184 range under multidimensional scaling, reflecting the fine-grained organization of upper extremity motor representations accessible with high-density surface sampling. Further analyses may be required to disambiguate whether these clusters reflect somatotopic organization as opposed to kinematic differences. In either case, however, the consistent and non-random population structure reflects information content relevant for decoding cortical signals, as evidenced by the ability of spatiotemporal features derived from high-gamma activity to discriminate movements using simple classifiers ^60^. For example, during a 4-way center-out joystick movement task (Fig. 4i-j), a convolutional neural network was able to decode with accuracy of 83% (chance = 25%), and precision and recall both at 83% (Fig. 4k-l), trained on 200 trials and tested on 47 hold-out trials.

### 5. Language-Related Responses in the Ventrolateral and Motor Cortex

High-density arrays (N=16) were implanted over the ventrolateral frontal and temporal cortices (Fig. 5a, c, f), including the inferior frontal gyrus (BA44), ventral precentral gyrus (BA4,6), and superior temporal gyrus (BA22, 42), in 16 subjects, yielding 12.2 hours of annotated awake recordings. Recordings were obtained during clinically standard language-mapping paradigms ^18^, including picture naming ^61^, counting, and word or sentence repetition. When clinically indicated, language-related regions were additionally localized using speech arrest ^18^ mapping prior to array placement.

During speech repetition tasks, subjects produced utterances drawn from vocabularies ranging from small (as small as 3 words) to larger lexicons (up to 50 words). Time-locked neural responses exhibited robust task-related modulation, with localized high-gamma (70–200 Hz) increases and concurrent low-frequency changes distributed across subsets of electrodes within each array. These responses were spatially heterogeneous at the sub-millimeter scale, revealing discrete clusters of electrodes with distinct temporal tuning profiles during word articulation ^62^ ^63^ ^30^.

With arrays placed over area 6v and regions identified as critical to speech (associated with speech arrest when stimulated) (Fig. 5f), the neural signatures during utterances of different words, when dimensionally reduced through locally linear embedding ^30^ ^64^ ^65^, form natural clusters in embedding space, presented via a t-distributed stochastic neighbor embedding (t-SNE) (Fig. 5d). These lower-dimensional features form a foundation from which classification at 92% accuracy (chance = 25%) (Fig. 5e) can be achieved with only 18 training samples, while supporting end-to-end detection (100% precision and 97% recall) and classification (79% accuracy on recalled instances), for a vocabulary size of three words and silence.

Decoding spoken words from face motor cortex (Fig. 5f) was more challenging than hand gestures but remained feasible. Dimensionality reduction via t-SNE visualizes words forming separable clusters in neural feature space (Fig. 5g), consistent with distinct articulatory representations at the electrode scale. Using multi-band spectral features with leave-one-out cross-validation, four-word classification accuracy reached 92% overall (per-class range: 85–100%; chance = 20%).

### 6. Facial Expression Responses in the Ventrolateral Cortex

In addition to speech production, some of the arrays (Fig. 6a-b) captured neural activity associated with non-speech facial expressions, including three to eight different expressions ^66^. The experimental paradigm alternated between a neutral resting state and targeted emotive expressions (Fig. 6c), allowing for the isolation of expression-specific neural signatures.

The recordings revealed that distinct facial expressions ^67^ elicit spatially separable patterns of high-gamma activity across the electrode array. As shown in the spatial maps, the facial expressions classically associated with certain emotional states (including disgust, fear, sadness, and others) activated unique, albeit partially overlapping, subsets of electrodes (Fig. 6d). These distinct spatial footprints enabled the classification of specific facial expressions using neural decoding models. A confusion matrix of the decoding performance across eight distinct facial motor tasks demonstrates non-chance classification accuracy for distinct expressions, with true positive rates reaching up to 67% for specific classes (including “shocked” and “grimacing” expressions) (Fig. 6e). Through the proxy of a cued facial posture paradigm, these results support that high-resolution surface electrophysiology may resolve the fine-scale cortical dynamics necessary to differentiate among complex emotive motor behaviors.

### 7. High-Resolution Tonotopy and Auditory Responses in Auditory-Associated Temporal Cortex

To investigate the micro-scale organization of auditory-associated areas, high-density arrays were deployed over the superior temporal gyrus (BA22, 42) in 3 subjects (Fig. 7a–b). These placements yielded over 3,000 recording channels of electrophysiologic data collected during anesthetized states, utilizing paradigms designed to isolate frequency-specific processing ^68^ ^69^ ^70^.

During passive listening tasks, neural activity was recorded in response to spectrally distinct acoustic stimuli, including pure tones and broadband pink noise, each presented with 50 trials (Fig. 7c). The high spatial resolution of the array revealed tonotopic organization within the superior temporal gyrus that is often spatially smeared in standard cortical recordings. As illustrated in the frequency-specific response maps, lower-frequency stimuli (e.g., 225 Hz tones) elicited maximal power in the 9.8–34.2 Hz frequency band in localized clusters of electrodes spanning approximately 10 mm^2^. Trial-averaged responses revealed that higher-frequency stimuli (e.g., 2773 Hz tones) only partially engaged the populations within the same cortical territory, while activating additional adjacent populations. In contrast, presentation of broadband pink noise recruited a broader distribution of electrodes, reflecting the wider spectral content of the stimulus. However, some subregions tuned to tonal responses were not responsive to pink noise. The spatial extents of 225 Hz, 2773 Hz, and pink noise were 10.4±0.3 mm^2^, 6.0±0.2 mm^2^, and 6.3±0.4 mm^2^, respectively, representing a pairwise centroid-to-centroid distance of 2.3±0.2 mm, 6.5±0.3 mm, and 4.3±0.3 mm for 225 Hz to 2773 Hz, 225 Hz to pink noise, and 2773 Hz to pink noise, respectively. These results demonstrate that high-density surface arrays can resolve the fine-grained spatial gradients of human tonotopy, capturing discrete frequency representations at the sub-millimeter scale.

### 8. Brainstem and Spinal Cord Surface Electrophysiology

The atlas extends high-resolution electrophysiological mapping beyond the cerebral cortex to include brainstem and spinal cord surface recordings. High-density arrays were successfully deployed over the cervicomedullary junction, ventral pontine surface, dorsal columns, and thoracic spinal cord, yielding a total of 3.3 hours of annotated recordings across 9 subjects (Fig. 8a–d). The flexibility of the thin-film arrays allowed conformal coverage of the recording regions, offering mapping targets less accessible to conventional electrodes. Recordings were obtained during clinically indicated procedures and included evoked responses time-locked to peripheral somatosensory stimulation and cranial nerve activation.

In the brainstem, arrays recorded evoked responses during trigeminal nerve stimulation ^71^, including selective activation of the ophthalmic (V1) and maxillary (V2) branches (Fig. 8e-g). Distinct spatial distributions of evoked signal amplitude were observed for V1 versus V2 stimulation, revealing branch-specific organization of trigeminal afferent activity on the exposed neural surface at the trigeminal root entry zone. Responses typically exhibited an initial peak at 3 ms post-stimulation. In cases with recordings obtained before and after nerve decompression, evoked responses were preserved and spatially localized across conditions.

Surface recordings over the posterior columns captured robust somatosensory evoked potentials with early latency components consistent with ascending dorsal column conduction (Fig. 8h-j). The initial positive peak response occurred at 13 ms after stimulation. The 400 µm electrode pitch enabled spatial discretization of these responses across the surface of the cervicomedullary junction, resolving gradients and propagation patterns that are typically observed as spatially averaged far-field potentials in standard intraoperative monitoring.

Recordings from the thoracic dorsal spinal cord (T4–T5) demonstrated high-fidelity, localized SSEPs elicited by left and right tibial nerve stimulation (Fig. 8k-m) ^26^, with peak response identified at 32 ms post-stimulation. Evoked activity exhibited clear lateralization, with spatially distinct clusters corresponding to ipsilateral sensory input. These recordings provide a direct characterization of the spatial structure of spinal surface potentials at sub-millimeter resolution.

Together, these data establish the feasibility of high-density surface electrophysiology in brainstem and spinal cord regions and provide an initial quantitative reference for evoked neural activity across surgically accessible neuraxial structures.

## Discussion

In this work, we present an initial functional atlas of the human central nervous system derived from high-resolution surface electrophysiology across 70 patients. By aggregating dense recordings spanning cortex, brainstem, and spinal cord, the atlas demonstrates that conformable, high-density surface interfaces can (1) resolve somatotopic organization at submillimeter spatial sampling density across multiple cortical functional regions ^50^ ^56^, (2) capture speech- and language-related dynamics from ventrolateral cortical fields ^30^ ^65^, (3) decode nuanced facial movements related to communication and the expression of emotion ^66^, and (4) map stimulation-evoked signatures from surgically accessible brainstem and spinal cord surfaces ^23^ ^26^. Collectively, these recordings have characterized reproducible functional organization and population dynamics at spatial and temporal scales that have not been accessible in prior human datasets. Rather than treating intracranial electrophysiology as a collection of isolated case studies, this work establishes a framework for systematic, cross-subject synthesis of human neural activity across multiple functional domains and bridges a gap in resolution between invasive penetrating arrays and noninvasive modalities.

Across the cerebral cortex, the atlas reveals fine-scale structure that is largely obscured by standard clinical recordings ^21^ ^56^. Submillimeter sampling resolves fine, two-dimensional transitions at canonical boundaries such as the central sulcus, including phase reversals and digit-level somatotopy, and captures spatially focal spectral modulations associated with movement, speech, non-verbal expression, and auditory processing. For instance, across subjects, digit representations generally followed the lateral-to-medial ordering within S1 expected on the basis of the classically described sensory homunculus ^50^ ^72^, while revealing partial overlap and fragmentation at the sub-millimeter scale that is not resolved by conventional centimeter-scale electrocorticography.

These observations are largely consistent with classical functional maps, but extend them by demonstrating that micro-scale cortical organization appears at least partially conserved across individuals when recordings are aligned in a common anatomical framework. This level of resolution enables mechanistic interrogation of population dynamics underlying motor control, speech articulation, and sensory processing, and provides features that are directly relevant for neural decoding and closed-loop neuroprosthetic applications.

A distinctive aspect of the atlas is its extension beyond the cerebral cortex. High-resolution surface recordings from the brainstem and spinal cord demonstrate that these regions—which are critical to sensorimotor function ^24^ ^26^—exhibit structured, spatially organized electrophysiologic signatures that can be captured with surface-based approaches. Recordings from the dorsal column pathway, trigeminal root entry zone, and thoracic spinal cord surfaces resolve stimulus-locked activity with sufficient fidelity to localize ascending sensory tracts and lateralization. Although sampling in these regions is necessarily opportunistic and anatomically constrained ^71^ ^73^, the results establish the feasibility of incorporating brainstem and spinal electrophysiology into a unified human atlas, and highlight a largely unexplored frontier for human systems neuroscience.

By occupying an intermediate regime between noninvasive imaging and penetrating microelectrode recordings, high-density surface electrophysiology offers complementary advantages. Compared with functional MRI, MEG, and EEG, surface electrophysiology provides direct measurements of neural population activity with millisecond precision and spatial resolution sufficient to resolve mesoscale organization ^1^ ^2^ ^3^. Compared with penetrating arrays, surface interfaces minimize tissue disruption and can be deployed transiently across a wider range of anatomical targets in standard surgical settings, enabling repeated recordings across patients to build a common anatomical map of functional cortical anatomy at high-resolution ^16^ ^17^ ^60^. High-density surface electrocorticography therefore fills a critical gap in the human neuroscience toolkit, enabling measurements that are simultaneously high-resolution, stable, and clinically practical.

The atlas is also designed as a resource for computational and translational work. Dense spatial sampling preserves high–spatial-frequency features that are often attenuated in lower-resolution recordings and that are increasingly important for modern decoding and machine learning approaches. Demonstrations of real-time motor ^15^ ^32^ ^36^ and speech ^30^ ^65^ decoding illustrate how these features can be exploited, but the broader value lies in enabling systematic benchmarking of models, cross-task generalization, and integration with structural and functional imaging. As the dataset grows, it may support the development of shared representational spaces for human electrophysiology. The dense multimodal dataset is well suited for modern machine learning approaches (including foundation models for neural decoding), cross-modal integration with structural and functional MRI, and computational investigations into micro-scale organization of the cortex, brainstem, and spinal cord.

Several limitations should be acknowledged. The dataset presented here reflects recordings obtained in heterogeneous clinical contexts with variable anatomical exposure and implantation duration, and sampling density is uneven across regions, particularly in the brainstem and spinal cord. In some cases normal anatomy was distorted by the presence of mass lesions. Tasks were necessarily adapted to clinical constraints and were not uniformly administered across all subjects; techniques for neural decoding required corresponding adaptations. Methods of anatomical localization were also dependent on clinical constraints, with array positions determined by available combinations of imaging, neuronavigation, and visual registration with anatomic landmarks. These factors limit the strength of some cross-region and cross-task comparisons in the present release. Nonetheless, by standardizing preprocessing, metadata, and coordinate registration, the atlas establishes a foundation upon which future data can be layered in a principled manner.

Looking forward, expansion of the atlas through further multi-site collaboration, standardized task batteries, and increased subject numbers will improve coverage and statistical power. Integration with chronic implants, where appropriate and feasible with respect to ethical and regulatory considerations, may enable longitudinal analyses of stability and plasticity. Coupling high-resolution surface electrophysiology with advanced computational approaches and multimodal imaging promises to refine our understanding of how distributed neural circuits give rise to human behavior and cognition. With appropriate cross-subject alignment techniques, a detailed map of the human central nervous system can gradually be constructed through the aggregation of functionally annotated data from multiple patients. From that viewpoint, this work reframes human electrophysiology as an atlas-building modality—one capable of bridging basic neuroscience, neurotechnology, and precision neurosurgery through a shared, extensible reference, to provide insight into human neural function.

## Methods

### Regulatory Approval and Ethics

All procedures were conducted under protocols approved by the Institutional Review Boards of the participating institutions. Written informed consent was obtained from all participants for temporary intraoperative or short-term postoperative research recordings and for data sharing in de-identified form. Device use complied with applicable investigational approvals under which the implants were performed. The device technology and prior human feasibility studies have been described previously ^15^. No unexpected adverse device effects were observed.

### Device and Array Design

Recordings were obtained using a thin-film, conformable microelectrode array ^15^ comprising up to 1,024 microelectrodes embedded in a polymer substrate ^74^. In selected cases, multiple arrays were used simultaneously. Array geometry, including electrode pitch and layout, varied slightly by device generation and surgical requirements. The nominal electrode pitch was 400 µm. Arrays were designed to maximize coverage of exposed neural surfaces while maintaining mechanical flexibility and conformability. Electrode materials, contact sizes, and impedance characteristics have been described in prior technical documentation and publications ^15^ ^31^. The recording platform incorporated head-stage amplification, multiplexing, and high-bandwidth data acquisition ^75^, enabling simultaneous capture of broadband neural signals across spatial and temporal scales ^76^.

### Surgical Implantation and Recording Context

Arrays were placed temporarily on neocortical, brainstem, or spinal cord surfaces using surgical exposure determined by the primary clinical procedure. In most (N=58) cases, arrays were deployed intraoperatively for short-duration recordings; in a subset (N=12) of cases, arrays remained in place for limited postoperative monitoring periods (typically days), during which intermittent recordings were obtained. For brainstem and spinal cord recordings, arrays were positioned on surgically exposed dorsal or ventral surfaces when clinically appropriate and consistent with safe surgical practice. All research arrays were removed according to protocol at the conclusion of the recording period. Additional details of the device hardware and surgical placement of the array are discussed in prior publications ^15^.

### Participants and Recording Targets

A total of 70 patients undergoing neurosurgical procedures that permitted safe temporary placement of thin-film electrode arrays were enrolled between 19 April 2023 and 9 January 2026. Informed consent was obtained preoperatively. A total of 86 arrays were deployed across the 70 patients in the final cohort, with 56 patients receiving single arrays and 14 patients receiving multiple simultaneous arrays for broader anatomical coverage. Together with 15 arrays being placed at 1 or more locations, this represented 102 total target locations. Per-subject recording targets are summarized in Supplementary Table 1. Recording targets included sensorimotor cortex (58 patients), including primary motor cortex (precentral gyrus), primary somatosensory cortex (postcentral gyrus), and the central sulcal region spanning both banks; frontal cortex (9 patients), including lateral prefrontal regions, ventrolateral frontal cortex including pars opercularis and pars triangularis of the inferior frontal gyrus; temporal cortex (7 patients), including superior temporal gyrus and superior temporal sulcus; and noncortical structures (9 patients), including brainstem (trigeminal nerves, cervicomedullary junction, and ventral pons) and spinal cord (dorsal columns at cervical and thoracic levels) surfaces.

Target selection was determined by the clinical surgical approach and anatomical exposure required for the primary procedure. Tractography was incorporated into preoperative navigation to assist with the verification of motor cortex location for relevant cases. Two methods were used to estimate electrode registration: CT-based localization of radiopaque markers (N=24) and intraoperative optical imaging paired with expert anatomical alignment (N=46). Each registration was verified to lie on the correct target gyrus, and registration error was estimated to be under 2 mm, which corresponds to the tolerance of the intraoperative image guidance systems used to assist in array localization.

Array implantation duration varied by clinical context. In 58 cases, arrays were placed and removed within a single operative session (median recording duration 22 minutes, range 3–66 min). In 12 cases, arrays remained in place postoperatively for extended monitoring over multiple sessions across multiple days (median total recording duration 4.5 h, range 0.9–52 h), during which recordings were obtained while patients were awake and able to participate in behavioral tasks. All arrays were removed at the conclusion of the recording period according to protocol.

Per-subject details including patient demographics, surgical indication, anatomical targets, recording duration, anesthetic state, and completed experimental paradigms are provided in a comprehensive fashion in Supplementary Table 1. Several figures show subsets of the total collection of deployed electrode arrays to maintain visual clarity.

### Peripheral Sensory Stimulation

Somatosensory evoked potentials were recorded during peripheral nerve stimulation performed by clinical intraoperative neuromonitoring teams under general anesthesia ^76^ ^77^. Stimulus timing was synchronized with neural recordings via TTL pulses for millisecond-precision temporal alignment. Electrical stimulation was delivered via needle electrodes with typical parameters ^78^: 1-3 Hz repetition rate, 200–500 μs pulse width, and 14–60 mA intensity (adjusted to 2–3 times the sensory threshold). Each paradigm included 30–200 stimulus repetitions for trial averaging.

#### Nerve-Level Stimulation

Median nerve stimulation at the wrist (N=22 patients) was recorded with arrays across the central sulcus to capture N20/P20 phase reversals ^27^. Tibial nerve stimulation at the ankle ^79^ (N=7 patients) was recorded over spinal or brainstem dorsal columns to characterize ascending somatosensory pathways ^26^. Trigeminal branch stimulation ^78^ (ophthalmic, maxillary, and mandibular; N=3 patients) was performed before and after microvascular decompression ^71^ with arrays over the root entry zone.

#### Digit-Specific Stimulation

Individual fingers were stimulated using metal ring electrodes (N=6 patients) to map fine-scale somatotopy within primary somatosensory cortex ^50^. Similar to electrical stimulation via needle electrodes, stimulation was delivered at 0.7-5 Hz repetition rate, 200–500 μs pulse width, and 14–60 mA intensity (adjusted to 2–3 times the sensory threshold), sufficient to achieve supramaximal activation during median nerve stimulation. Each paradigm included >100 stimulus repetitions per digit for trial averaging.

#### Vibrotactile Stimulation

In N=13 cases, vibrating motors delivered 0.5 s pulses at 150 Hz to fingertips (N=6) or facial (N=7) regions ^52^ (15–30 repetitions per block) as an alternative to electrical stimulation. Timing was controlled by an Arduino microcontroller with TTL synchronization.

### Cued Hand and Face Motor Tasks

Awake motor tasks were performed in two contexts: intraoperatively during awake craniotomy ^18^ with functional mapping (N=18 patients), and postoperatively in the intensive care unit or neurosurgical ward (N=9 patients). All tasks were presented on a display screen positioned approximately 60 cm from the patient at eye level. Task timing and behavioral data were synchronized with neural recordings via digital trigger pulses and timestamps.

#### Hand Gesture Tasks

In N=21 patients, discrete hand and finger movements were cued using visual prompts displayed on a screen. Each trial began with a 1-2 s rest period, followed by presentation of a gesture cue (image or text instruction) for 2-3 s. Patients were instructed to execute the cued gesture and hold it for 2-3 s until a return-to-rest cue appeared. Gesture sets varied by patient and clinical context but typically included individual finger flexion and extension (thumb, index, middle, ring, little), fist closure, hand opening, and wrist movements. Each gesture type was repeated 15-60 times. In all cases, the timing of the task prompts relative to neural data for motion and rest were logged. In N=19 cases, hand kinematics were recorded using motion capture gloves, including StretchSense MoCap Pro with capacitive sensors at each finger joint (Auckland, New Zealand), Manus Metagloves Pro with electromagnetic field (EMF) tracking of fingertips (Geldrop, Netherlands), and Movella Xsens Awinda sensors with inertial measurement units (Enschede, Netherlands). StretchSense glove data were sampled at 100 Hz and time-aligned with neural recordings via serial port triggers, while Manus and Xsens data were aligned in software using the computer clock. In N=14 cases, video recordings (30-60 fps) were also collected for offline movement annotation.

#### Cursor Control Tasks

In N=14 patients, continuous hand movements were recorded during center-out reaching tasks. Patients used a 2-axis joystick ^80^ (Logitech G X56 “HOTAS”) or game controller (Sony PlayStation) to control a cursor on a screen to reach a highlighted target. In some cases, the cursor was moved from a central starting position to one of 4-8 radially arranged targets (target diameter 4.5 cm, target distance 9.5 cm from center to center of the target in the x-direction and 5.5 cm from center to the center of the target in the y-direction). Each trial began with the cursor held at center (1-2 s), followed by target appearance and a “go” cue. Each target direction was repeated 15-30 times in pseudorandomized order (75 x-y trials per session). In other cases, the patient was presented with a WebGrid-like target-chasing task ^60^ with grids sized with 4×4, 8×8, or 12×12 squares and moved the cursor to a highlighted square. Once the cursor reached that target, another square was highlighted in a randomized grid position. In all cases, patients were instructed to reach the target as quickly and accurately as possible. Successful trials required cursor entry into and hovering within the target zone for 0.3–1 s. Joystick position was sampled at 100 Hz and time-synchronized with neural data. Cursor position, velocity, and trial event markers (target onset, movement onset, target acquisition) were logged for offline analysis.

#### Facial Expression Tasks

In N=8 patients with arrays positioned over ventral precentral cortex or inferior frontal regions, facial expressions were cued using emoticon images and text labels displayed on the screen. Each trial began with a 1-2 s neutral face baseline, followed by an expression cue (1-2 s presentation), during which patients were instructed to produce and hold the cued expression until a return-to-neutral cue appeared (blank screen). Expression sets varied but typically included a neutral (relaxed) expression as well as 3-8 distinct facial expressions drawn from: “Angry,” “Smirking,” “Frustrated,” “Terrified,” “Happy,” “Sad,” “Shocked,” and “Neutral”; as well as “Grimacing,” “Lip Pucker,” and “Tongue Protrusion.” Each expression was repeated 15-30 times in pseudorandomized blocks. Video recordings of the face were acquired, with appropriate consent, using a digital camera (resolution 1920×1080 or higher, frame rate 30-60 fps) positioned to capture full facial view. Video was synchronized with neural recordings via LED flash triggers visible in the video frame at trial onset, enabling frame-accurate alignment for offline analysis.

#### Data Synchronization

All behavioral data streams (motion capture, joystick, video timestamps, EMG) were synchronized with neural recordings using one or more of the following methods: TTL trigger pulses sent from the stimulus presentation computer to digital input channels on the recording system, serial port communication with millisecond-precision timestamps, or visible or audible markers recorded simultaneously in behavioral and neural data streams via analog ports. Synchronization accuracy was verified post-hoc by aligning known event markers, achieving consistent alignment precision <10 ms.

Per-subject task completion details, trial counts, and recording durations are provided in Supplementary Table 1.

### Speech and Language Tasks

Speech tasks were performed in N=21 patients during awake recordings. Speech was recorded via directional microphone ^30^ (44.1 kHz, synchronized with neural data) and annotated for phonetic content. Standardized task batteries ^61^ were used where possible to allow cross-subject aggregation. The paradigms were divided into clinical tasks used for language mapping during awake neurosurgery, and research-specific tasks designed to isolate the neural dynamics of speech production and perception.

#### Clinical Mapping Tasks

Clinical mapping tasks were performed in N=4 patients. During awake language mapping, these patients performed object naming using standardized line drawings, serial counting, and cued speech tasks including sentence completion and repetition adapted from standard aphasia batteries. These tasks were selected to probe lexical retrieval, automatic speech, and phonologic processing during cortical stimulation. Stimuli were presented using a tablet-based digital interface with standardized timing and rapid task switching, enabling consistent stimulus delivery and real-time response monitoring. Patient responses were assessed intraoperatively for speech arrest, anomia, and paraphasic errors to localize eloquent language cortex.

#### Cued Word Reading

Visually cued word repetition tasks were performed in N=13 patients. To investigate the transition from semantic processing to motor articulation, patients were asked to read 3-4 words from a screen with delayed articulation (a read-wait-speak paradigm) ^83^. Word sets were designed with specific phonetic properties. One minimal phoneme set ^30^, {“ark”, “car”, “rock”}, exhibited high phoneme reuse. Another, {“teeth”, “key”, “wow”, “mom”}, exhibits substantial contrast (minimal phonemic overlap). These sets were cued alongside a designated silence condition. To temporally separate reading from articulation, the patient was asked to wait between word presentation and word repetition, cued by a color change on the screen. Each trial commenced with a blank screen for 1.0 to 1.5 s. A red “preparation” screen then displayed the target word for 1 to 2 seconds, during which the patient was instructed not to speak. This was followed by a green “cue” screen, giving the patient a 5 s window to speak the word aloud.

#### Sentence Repetition

To capture the complex kinematic and phonetic features of continuous speech, N=4 patients repeated sentences either played aloud through a speaker, read aloud to the patient, or displayed on a screen for the patient to read. For this task, we utilized the 50-phrase-AAC set, which contained 50 sentences composed of a vocabulary of 119 unique words ^65^, or a subset of sentences from the TIMIT Acoustic-Phonetic Continuous Speech Corpus ^84^.

Tasks aligned to triggers were recorded with synchronized behavioral logs. To ensure precise, millisecond-level alignment for subsequent analysis, the audio was co-recorded directly with the neural data.

### Auditory Tasks

Passive auditory tasks were performed on N=3 patients with arrays over superior temporal gyrus. The patients were presented a pure tone (225 or 2773 Hz) or pink noise at 2 Hz, with over 100 trials. For one patient, the first six principal tones (from middle C to A) in C major were presented, followed by 3 repetitions of a piece of music composed of the six tones.

### Recording and Preprocessing

#### Data Acquisition

Signals were sampled at 20 kHz ^76^ using either an Intan RHD recording system ^75^ with 16-bit analog-to-digital conversion or custom electronics. Reference electrodes were either placed at the Fz location using a needle electrode, or placed subdermally or epidurally near the edge of the surgical exposure via a needle or gold cup electrode. The ground electrode was placed either at the shoulder using a needle electrode, or at a skin patch located far from the surgical site. Electrode impedances were measured at 1 kHz; electrodes >4 MΩ were excluded (N=77 arrays and N=69,254 included electrodes, mean 1.17±0.50 MΩ, 89.1±12.2% yield rate).

In early device iterations (N=5), 982 of the 1,024 microelectrodes were routed to signal amplifiers. While the remaining 42 contacts were in each case successfully placed in interface with the cortical surface, these specific channels were not sampled for electrophysiologic recordings or impedance data. In a subsequent configuration of the custom electronics (N=8 arrays across N=7 patients), 960 electrodes were dedicated to the recording pathways, with the remaining 64 electrodes specifically routed to hardware optimized for impedance characterization.

#### Preprocessing

Common average referencing ^81^ was applied to cortical arrays on a case-by-case basis depending on whether it improved decoding accuracy. Otherwise, no re-referencing was performed. Line noise was filtered out (60 Hz notch with harmonics). In analysis-specific filtering, high-gamma was defined as 70-200 Hz ^57^, and beta was defined as 12-30 Hz ^85^. Zero-phase finite impulse response filters ^86^ were used throughout.

### Localization and Cross-Subject Registration

#### Cortical Arrays

Electrode positions were localized using intraoperative photography, post-implant CT or MRI (when available), and preoperative MRI (when available). Cortical surfaces were reconstructed from T1-weighted MRI using FreeSurfer (version 7.4.1) ^40^. In RAVE ^87^ ^88^, electrodes were registered and projected onto individual pial surfaces, then transformed to fsaverage template space via surface-based spherical registration. In most cases where only intraoperative photography and intraoperative image guidance (“neuronavigation”) were available, electrode positions were registered to an example patient brain, prior to transformation to the standard brain. Registration quality has a maximum landmark error of 2 mm, corresponding to the maximum accepted error of the intraoperative image guidance systems.

#### Brainstem and Spinal Cord Arrays

Array positions were determined with the assistance of intraoperative image guidance (“neuronavigation”) and from intraoperative photographs, and positions were verified with respect to nearby anatomic landmarks (trigeminal root entry zone, pontomedullary junction, and vertebral levels). Cross-subject alignment used rigid-body transformation of landmark positions due to limited detail in standard atlases. Coordinates were expressed in custom landmark-aligned template space.

#### Quality Control

All registrations were manually inspected. Sulcal alignments were verified for cortical arrays; landmark distances were measured for all regions.

#### Coordinate Formats

Electrode positions are provided in native space, fsaverage surface space, MNI152 volumetric space ^5^, and brainstem or spinal cord template space. Transformations and surface meshes are saved together with the underlying data.

### Statistical Analyses

All analyses were performed in Python or R. Statistical significance was assessed at α = 0.05, two-tailed, unless otherwise noted.

#### Time-Frequency Analysis

Power spectral densities were computed for 1 second prior to and following activity onset, using the raw signal downsampled to 2 kHz. The signal was further z-scored to a time window [-875, -625] milliseconds prior to activity onset. The Welch method was used to extract power spectral density (sampling frequency = 2 kHz, nperseg = 2048). The ratio between spectral power pre-activity and during-activity was then computed for each frequency step, as segmented during application of the Welch method, followed by a mean across the steps.

#### Decoding

Spectral power envelopes derived from filters applied over predetermined frequency ranges were used to construct model features. Raw data were first filtered for specific bands, then envelopes were computed via absolute value of the Hilbert transform ^89^. High-gamma band (70–150 Hz) was included in all decoding architectures, while other frequency bands were included on a case-by-case basis. These frequency bands were all between 0.5 and 200 Hz, segmented into smaller frequency ranges, such as, but not limited to, local motor potential (0.5–4 Hz), beta band (12–30 Hz), and a wider gamma band (70–200 Hz). Models included logistic regression, convolutional neural networks (CNN), recurrent neural networks (RNN), a combination of both (CRNN), long-short-term memory networks (LSTM), inception-inspired architectures, and other techniques. Evaluation was performed on stratified 5-fold cross-validation, with models trained on 80% of trial data, and tested on the remaining 20%. Decoding metrics included accuracy and correlation.

#### Cross-Subject Spatial Analyses

To quantify the spatial relationship between electrode arrays and the central sulcus, the Euclidean distance from each electrode to the anatomic sulcal boundary was computed in MNI152 space. The boundary was defined as the set of Destrieux atlas voxels where the precentral and postcentral gyrus labels are directly adjacent, including central sulcus-labeled voxels neighboring either gyrus; this yields a representation of the sulcal boundary at the cortical surface rather than its deep fundus. For each electrode, the distance to the nearest boundary voxel was computed via a k-d tree spatial index. Per-deployment summary statistics (mean, median, standard deviation) were reported.

#### Sample Size

The number of subjects available per region varied (1–25, Table 1), and this largely reflects the opportunistic nature of surgical access that contributed to cohort construction.

## Acknowledgments

We thank the study participants and their families. Several of the coauthors and collaborating institutions have received grant support for portions of their contributions to the work reported here: NSF BRAIN IUCRC 2310939 (West Virginia University Rockefeller Neuroscience Institute), NIH R01-NS125026 (Northwestern University Feinberg School of Medicine), NIH K12-NS129164 and R01-EY035826 (Perelman School of Medicine at the University of Pennsylvania), NIH R01-NS134410 (Massachusetts General Hospital), the Weill Neurohub Clinician-Scientist Award (University of Washington), and the Fitzgerald Translational Neuroscience Fund (Thomas Jefferson University). We also thank the intraoperative and extraoperative clinical teams at each institution for their contributions. We acknowledge the contributions from clinical research staff at the collaborating institutions who enabled the data collection and analysis reported here, including Molly Summers, Pat Dekeseredy, Tristen Cheuvront, Lizbeth Nunez, Katherine Belity, Emily Svendsen, Kevin Buttet, Kim Agosto, Marie Kerr, James Carter Mitchell, Domenic Annand, Barshana Karki, Brian Hsueh, Yangling Chou, Brian Coughlin, Amy Anderson, Miles Mahon, Danielle VanBrabant, Markey Olson, Ellen Carl, Nicole Armstrong, Alessandro Napoli, Phyo Thuta Aung, Jessica Rinaldi, Nico Agosti, Erica Jones, and Maclain Capron. We also acknowledge Jaime Chong (Precision Neuroscience) and Danielle VanBrabant (Barrow Neurological Institute) for their contributions to the medical illustration and graphic design content of manuscript.

## Data and Code Availability

All processed data, anonymized raw traces, electrode coordinates (in atlas space), behavioral logs, and analysis code have been carefully curated and archived. Data and code availability will be determined concurrent with publication of the final peer-reviewed version of this manuscript, and will be subject to legal and ethical constraints inherent to work with human subjects. Requests for data and code may be directed to the corresponding author.

## Supplementary Materials

**Supplemental Table 1:**
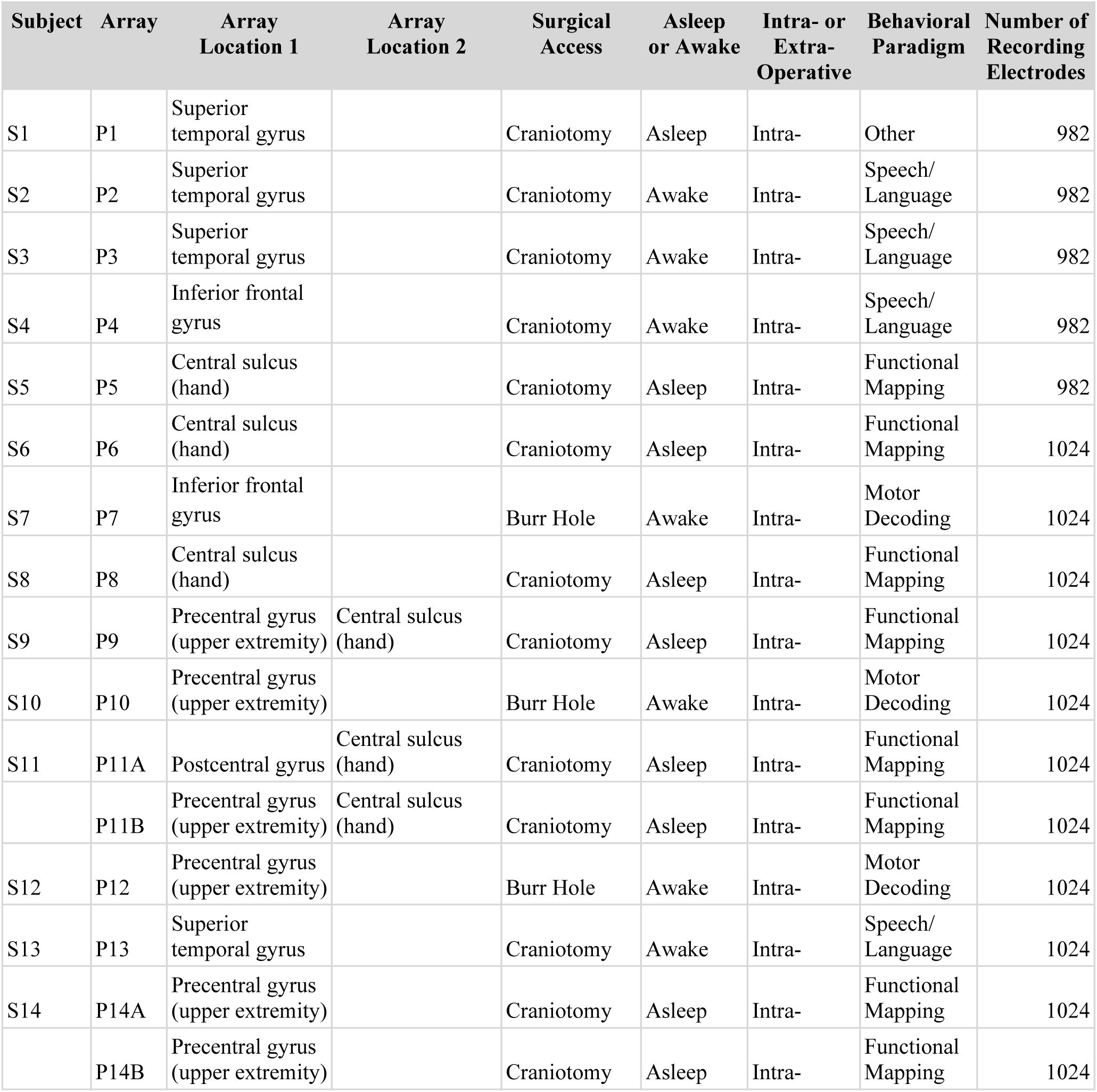
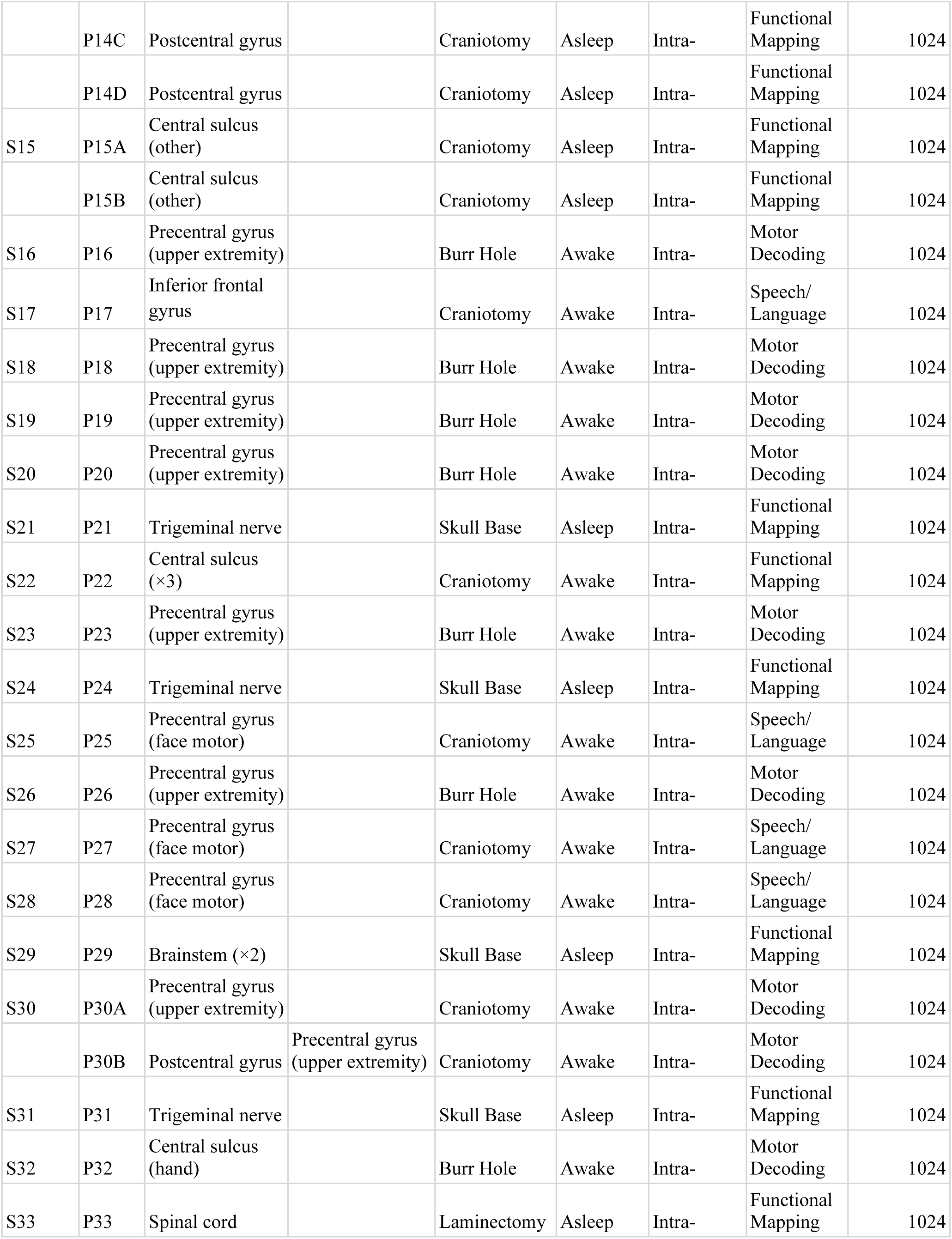
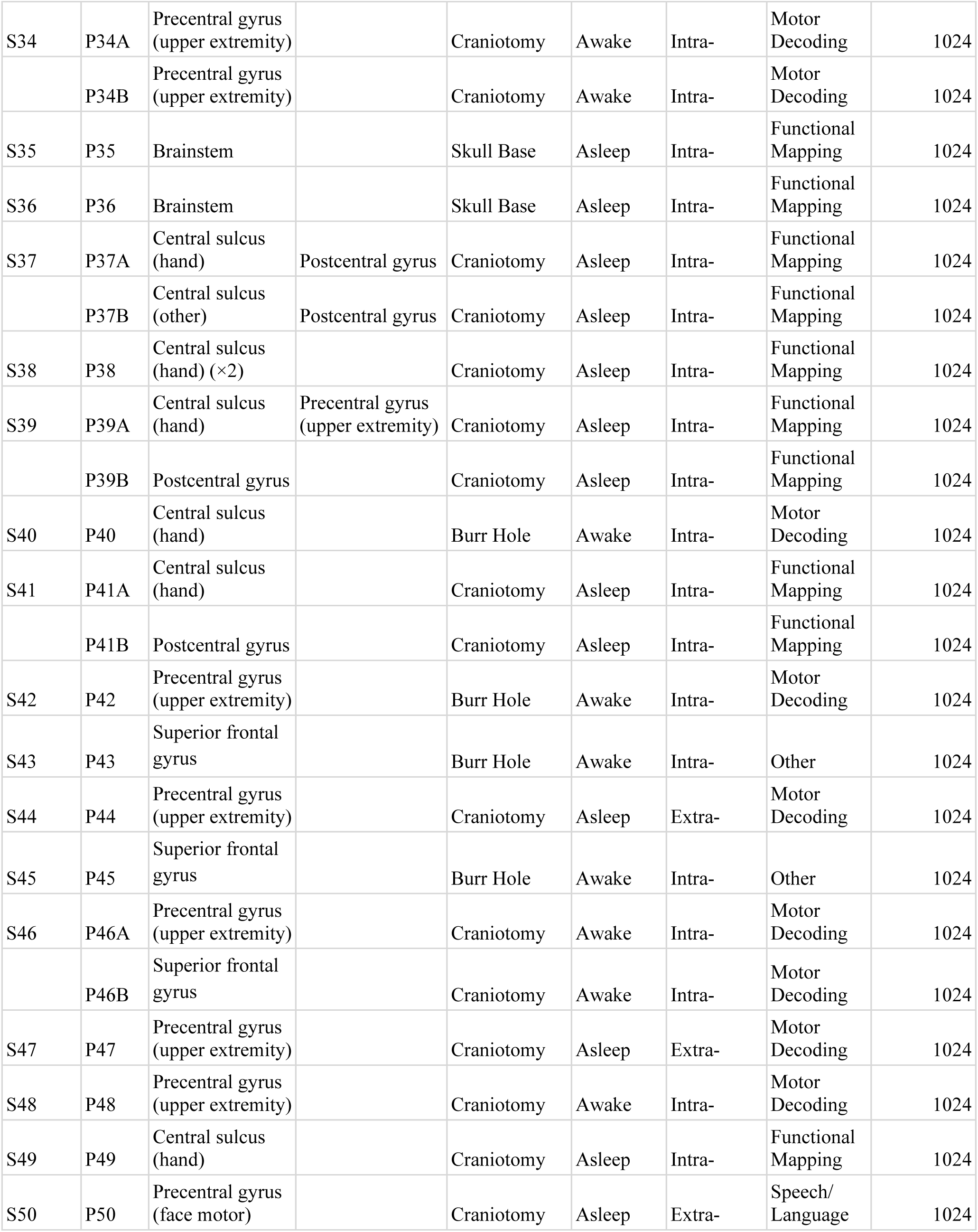
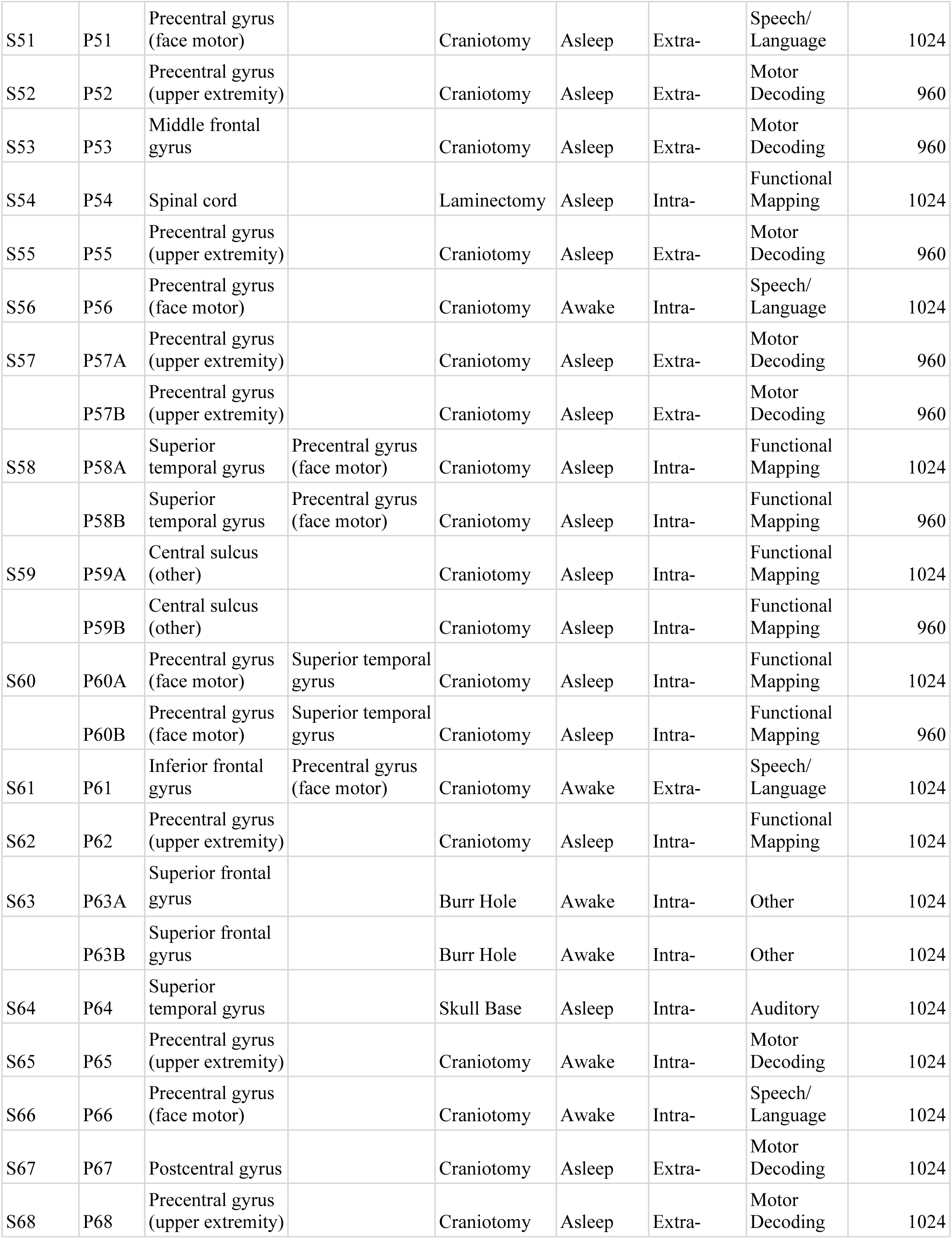
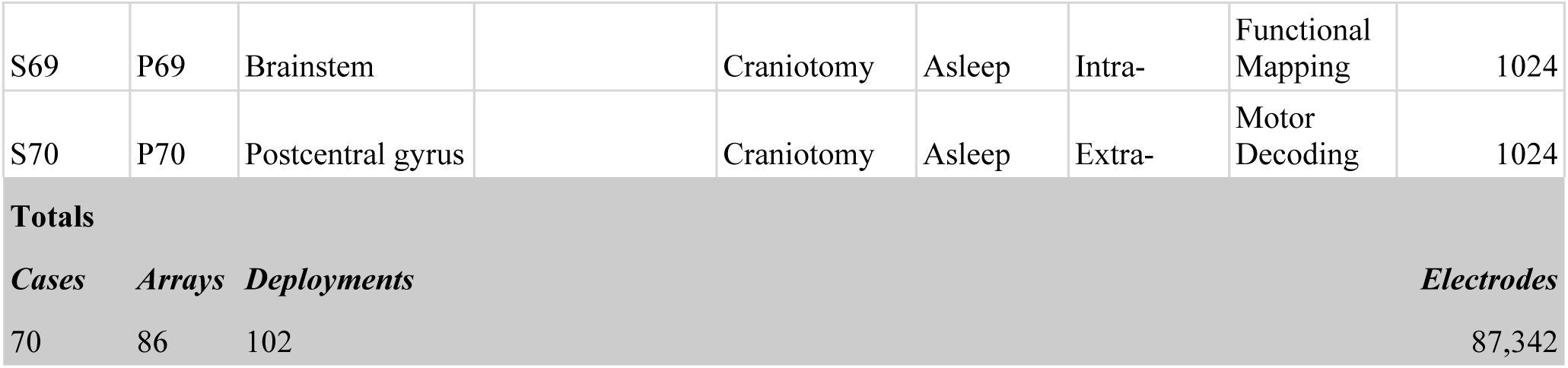
Details of Individual Cases Included in the Atlas. Array placement, surgical and experiment details for each subject (S-labels) implanted with one or more corresponding electrode arrays (P-labels). In some cases, two or more arrays were placed simultaneously in two locations and listed in the table. In addition, one array may have been placed over multiple locations during one intraoperative session. The “Awake or Asleep” category identifies whether the patient was awake or asleep during the intraoperative recording. The “Intra-” category refers to whether the recordings took place only during surgery (“Intraoperative”) or during post-surgical recovery in the hospital (“Extraoperative”).

**Supplemental Figure 1:**
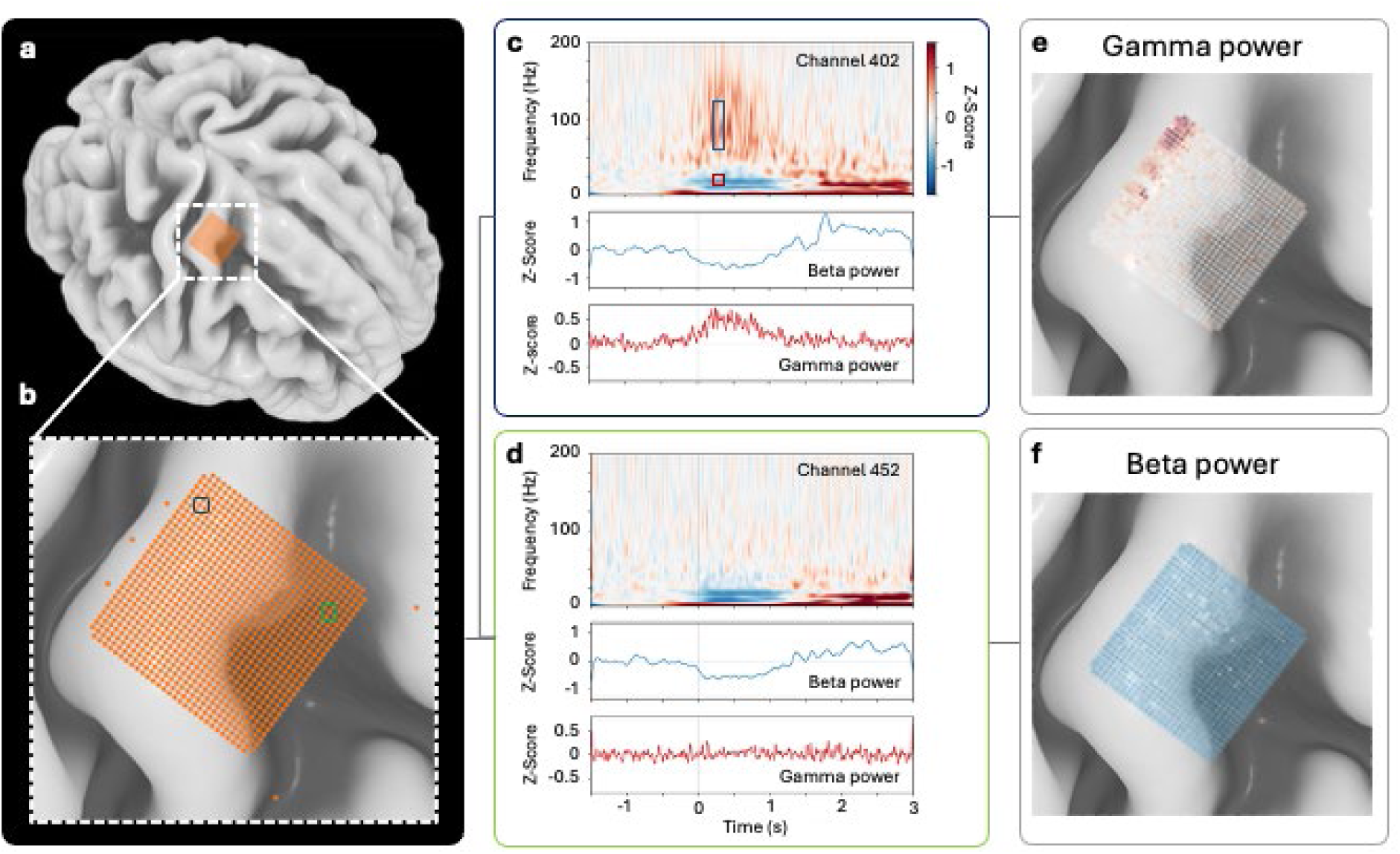
Example Event-Related Spectral Frequency (ERSF) and Spatial Distribution. (a) An array was placed over the hand knob area of the M1 primary motor cortex (BA4) on the right hemisphere. (b) Zoomed-in view of the array. The signals of the two boxed electrodes (dark blue and green) are presented in (c) and (d). (c, d) Event-related spectral frequency (ERSF) of two example electrodes, recorded over 61 finger movement trials. Kinematics data were measured using motion capture gloves. The first derivatives of positional measurements were used to compute finger velocities, and then thresholded to 4 standard deviations of baseline movement for the identification of movement onset, aligned at time zero. (e, f) Spatial distribution of gamma and beta activation 375 ms post-onset (red and blue boxes in panel (c), respectively), demonstrating classical beta desynchronization and gamma activation observed for the movements. The color scales are matched to the ERSFs in panels (c) and (d).

